# Electrophysiology of multisite direct electrical stimulation in the human brain

**DOI:** 10.64898/2026.11.11.765993

**Authors:** Devang Sehgal, Angelique C. Paulk, Peter N. Hadar, Jaquelin D. Peralta, Pariya Salami, John D. Rolston, R. Mark Richardson, Sydney S. Cash, Rina Zelmann

## Abstract

Human cognition and neurological disorders reflect properties of distributed networks, yet how best to engage these networks therapeutically remains unknown. Multisite stimulation is a principled strategy for engaging brain networks, but a key translational barrier is understanding how a network combines concurrent inputs from multiple nodes. Here, we investigated the electrophysiology of simultaneous multisite direct electrical stimulation in 20 participants implanted with stereo-EEG electrodes. Synchronized dual-site stimulation engaged a larger effective network than single-site stimulation. Linear superposition explained most responses, but distinct nonlinear responses were also observed. Sublinear responses, consistent with saturation, occurred proximal to stimulation sites, while supralinear responses arose in network-connected sites during the response’s late component. Moreover, low-intensity dual-site stimulation matched or exceeded single-site stimulation at higher intensities. These principles extended to three-site stimulation. These findings systematically characterize how responses to simultaneous multisite stimulation interact in the human brain, a foundational step towards principled, network-guided neurostimulation therapies.

## Introduction

Direct electrical stimulation (DES) of the brain is an effective therapeutic approach for a wide range of neurological^1,2,3^ and neuropsychiatric^4,5,6^ disorders, with over 250,000 people undergoing treatment worldwide^7^. Although brain function and disorders are increasingly understood as properties of distributed networks rather than single anatomical regions^8–10^, how best to engage these brain networks remains a central open question^11^. Treatment has been largely limited to stimulation delivered at a single anatomical target associated with the disease, or a presumed hub such as the thalamus^2,10^. Clinical use has shown variable efficacy across patients and diseases^3^, while patients often undergo frustrating trial-and-error approaches before experiencing therapeutic benefit^12^, a process which can last years^13^. With one node, therapeutic efficacy is pursued largely by raising stimulation intensity, which can lead to side effects, such as painful sensations, paresthesia, and disruption of sleep or cognition^14,15,16,17^. The mismatch between a network model of disease and single-node treatment constrains the efficacy that stimulation can reach. Stimulating several nodes at once offers the possibility of engaging a wider network while reducing the stimulation intensity required at any one location. However, a key gap in our understanding is how a network combines simultaneous inputs from multiple nodes - whether the responses simply add, or interact nonlinearly.

As an extension of the single-hub approach, clinical applications have used symmetric bilateral targets with deep brain stimulation (DBS) for Parkinson’s disease or independent hubs targeting multiple foci with responsive neurostimulation (RNS) for epilepsy. Recent reports have begun to explore multi-lead and multitarget DBS for multifocal^18^ and generalized^19^ epilepsy, and for Parkinson’s disease^20^, hinting at the possibility of managing comorbid disorders^21^ or combining targets in treatment refractory disease^22,23^. Recent computational^24^, pre-clinical^25,26^, and memory studies^27,28^ have also suggested the potential value of multisite stimulation. However, these strategies have been developed empirically, with each site selected and often programmed independently. Systematically characterizing the electrophysiology of simultaneous multisite stimulation in humans is the first step towards principled, physiology-based, network-targeted multisite strategies.

A natural first-order model of the response to simultaneous multisite stimulation is the linear superposition of the responses to each site alone^29^. Under this hypothesis, a second site could broaden the network engaged through the addition of weaker, sub-threshold responses, while interactions at sites effectively connected to both stimulation locations could cause responses to exceed or fall below the sum. Distinguishing these effects determines what a second stimulation site contributes and whether the sites can be programmed independently.

To test this hypothesis, we investigated the dynamics of multisite stimulation. We delivered single pulses of electrical stimulation (SPES) to one, two, or three sites simultaneously in participants implanted with depth electrodes and analyzed the evoked responses across the brain. Evoked responses, or cortico-cortical evoked potentials (CCEPs), have been used to map functional networks^30,31^, delineate epileptic areas^32^, and study physiology^33,34^. Since stimulation was delivered to both cortical and subcortical (mostly thalamic) regions, the term time-series evoked potential (TSEP) is used here instead. We compared the responses to dual-site stimulation with the corresponding single-site responses and their sum at three levels: changes in network engagement, deviations in the time series evoked activity, and the key features of the evoked response. This allowed us to quantify how well single-site responses predicted dual-site responses, and to characterize the spatial, temporal and intensity dependence of departures from linear superposition. Finally, we tested how well the understanding developed in the two-site case extends to three-site stimulation to assess the generalizability of multisite stimulation biophysics. Together, these measurements provide a systematic characterization of how responses to synchronous stimulation interact in the human brain across regions and stimulation intensities.

## Results

Synchronous dual-site SPES was delivered at 87 pairs of sites across 20 participants (3–6 stimulation pairs per participant) with drug-refractory epilepsy implanted with stereo-EEG (sEEG) electrodes for clinical monitoring (Supplementary Table 1). Single-site SPES was interleaved at every dual-site stimulation site. Stimulation was delivered at 7 mA in all participants (N = 20; n = 87 pairs), and in a subset at 4 mA (N = 17; n = 66) and 2 mA (N = 7; n = 27). Intracranial EEG (iEEG) was recorded from 2,561 bipolar channels across participants after exclusion of channels with epileptiform activity, seizure-onset-zone involvement or excessive artifact (Methods). Recording coverage was concentrated in frontal and temporal regions but also included parietal and occipital cortex, cingulate, and the thalamus (Supplementary Figure 1). Which pairs to stimulate was selected, when possible, following a screening SPES session to obtain recording sites where evoked responses to individual single-site stimulation converged and could potentially interact (Methods). For each stimulation pair, we labelled the two sites ‘A’ and ‘B’, the simultaneous stimulation ‘A&B’, and the hypothesized linear sum of single-site responses ‘A+B’.

### Dual-site stimulation increases network engagement across brain regions and at distant sites

To test whether dual-site stimulation increased network engagement, we compared the channels responsive to synchronized dual-site stimulation with the union of channels responsive to single-site stimulation of either location, and identified the additionally responsive channels, i.e. responsive to dual-site but not to either single-site stimulation (Figure 1a). Responsiveness was defined by detection of an evoked potential in the time-series data (TSEP criteria, Figure 1b). Across all stimulation pairs at 7 mA, 20.4 ± 1.3% channels were additionally responsive to dual-site stimulation (mean ± s.e.m., n = 86 pairs meeting criteria for inclusion (Methods), Figure 1c, examples in Figure 1d).

Analogously, to consider the effect of dual-site synchronized stimulation on eliciting local neuronal firing, we computed the High Frequency Activity (HFA criteria - gamma band power from 70 Hz to 150 Hz), since HFA can be a proxy for local neuronal firing^35,36^ (Figure 1b). We found 25.1 ± 2.7% channels additionally responsive by HFA criteria (n = 84, Figure 1c, examples in Figure 1e). Additional responsiveness is observed stably across a range of responsiveness thresholds (Supplementary Table 2).

A natural question is whether this additional responsiveness depends on the brain region stimulated. We compared the two anatomically defined groups of stimulation pairs most frequently present in our dataset: frontal, with both stimulation sites in the frontal lobe of the same hemisphere (n = 14), and thalamic involvement, with one or both sites in the thalamus (n = 14). Neither criterion reached significance (frontal vs thalamic involvement, two-sided permutation test; TSEP: mean difference +6.1%, p = 0.21, Cliff’s δ = 0.35; HFA: −9.5%, p = 0.33, δ = −0.06; Figure 1f). Thus, additional responsiveness appears relatively similar across the brain (rates for all stimulation regions in Supplementary Figure 2 and Supplementary Table 8).

To assess the effect of proximity to the stimulation sites, we computed the fraction of additionally responsive channels as a function of minimum distance from either stimulation location, pooling channels across all pairs (Figure 1g). Within 15 mm^37^ (which separates near-field local responses from propagated distant activity^38^) of the nearest stimulation site, few channels were additionally responsive (close to 4%), while >20% of sites were observed to be additionally responsive further than 35 mm. These results held for both TSEP and HFA criteria. Beyond 90 mm, fewer than 5 channels per bin responded to A or B. Additional responsiveness is predominantly distal to the stimulation sites, consistent with network-mediated engagement.

**Figure 1:**
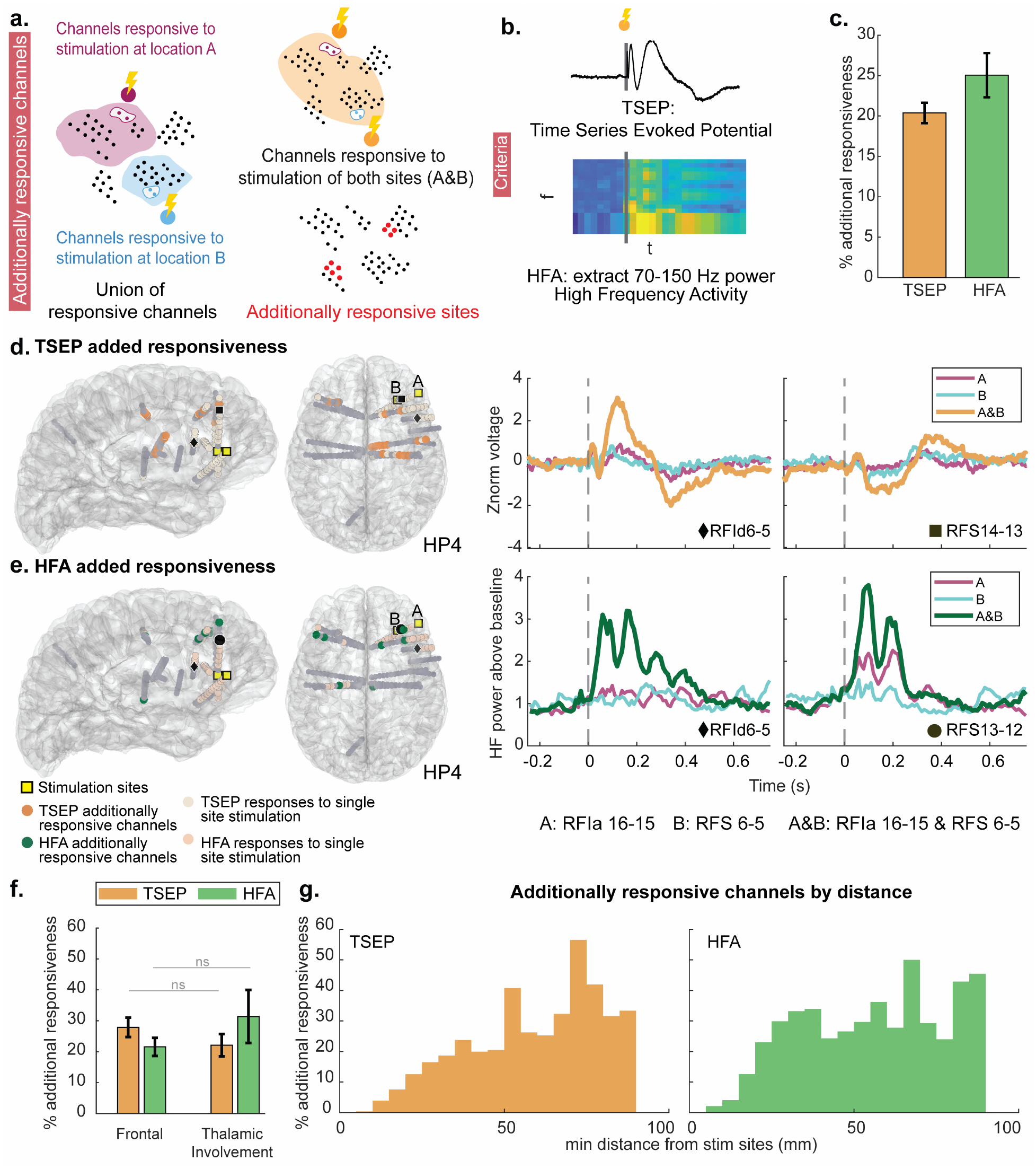
Effects of dual-site stimulation on network engagement. **a**, Additionally responsive channels are identified by comparing the channels responsive to simultaneous dual-site stimulation with the union of channels responsive to either single-site stimulation. Red dots mark additionally responsive channels. **b**, Responsiveness criteria. (1) A channel is considered TSEP responsive if an evoked potential is detected in the time-series activity. (2) A channel is considered responsive by the HFA criteria if an increase in high gamma power (70-150 Hz) is detected in response to stimulation. HFA is a correlate of local neuronal firing. Vertical gray lines mark stimulus onset. **c**, Percentage of channels additionally responsive to dual-site stimulation at 7 mA, relative to the union of channels responsive to either single site stimulation, by the TSEP (20.4 ± 1.3%) and HFA (25.1 ± 2.7%) criteria. Mean ± s.e.m. across stimulation pairs (n = 86 TSEP and 84 HFA pairs). **d**, TSEP additionally responsive channels on a glass brain (left) and example time-series responses (right) for one participant (HP4; stimulation of right pars orbitalis, RFIa16-15, and right lateral orbitofrontal cortex, RFS6-5). Pale dots mark channels responsive to single-site stimulation and gray dots other recording contacts. Black symbols mark the channels shown on the right. Traces show responses to single-site stimulation at A or B and to simultaneous dual-site stimulation (A&B). **e**, As in d, for the HFA criteria: HFA additionally responsive channels (left) and high-frequency power relative to baseline (right). **f**, Percentage of additionally responsive channels at 7 mA for the frontal and thalamic-involvement groups. Two-sided permutation test; n = 14 and 14 pairs; ns: not significant. **g**, Percentage of additionally responsive channels as a function of Euclidean distance from the nearest stimulation site, pooled across all 87 pairs at 7 mA. Bins with fewer than 5 channels responsive to A or B are not shown. Additionally responsive channels are observed more frequently distal to stimulation locations.

### Widespread linear superposition in response to dual-site stimulation

If dual-site stimulation broadens the overall network engaged, does it also change the physiology of any given response? We compared the response to dual-site stimulation (A&B) against the linear sum of the corresponding single-site responses (A+B), first asking whether A+B predicts whether a TSEP is observed. The hypothesized sum, i.e. ‘A+B’, was computed by adding the medians of the responses for ‘A’ and ‘B’ separately, and the TSEP detector was then run on this summed signal, with the same criteria as for recorded responses, to obtain the set of channels predicted to be responsive. Framed as a classification problem, a channel is correctly classified when A+B and A&B agree on its responsiveness, a false positive is a channel where A+B predicts a TSEP that is absent in A&B, and a false negative the converse (Figure 2a). Across all recording channels at 7 mA, A+B correctly classified 86.7% of channels, while 5.2% were false-positives and 8.2% false-negatives (Figure 2c).

### Distinct nonlinear responses across individuals

While a linear sum explained most responses, several channels showed distinct nonlinear responses that either exceeded the sum (supralinear) or fell below it (sublinear) (representative examples in Figure 2f). To identify these nonlinearities, we conducted cluster-based bootstrap tests between the time series responses of ‘A+B’ vs ‘A&B’. A channel was classified as supralinear (sublinear) if it exceeded (fell below) the sum for more than 75 ms (Figure 2b; other durations in Supplementary Table 3). A stimulation pair could produce linear, supralinear, and sublinear responses at different recording channels (Figure 2e-g).

Across all participants and stimulation pairs at 7 mA, among channels responsive to stimulation at both locations, i.e. effectively connected to both (20.6 ± 1.6 channels per pair; n = 63 pairs), 13.3 ± 2.1% were supralinear and 7.5 ± 1.4% sublinear (Figure 2d), with 1.6 ± 0.4% showing both substantial supralinear and sublinear segments. Among the larger set of channels responsive to either single site, A or B (56.4 ± 3.1 channels per pair; n = 86), the rates fell to 5.9 ± 1.0% supralinear and 3.8 ± 0.6% sublinear, indicating that nonlinearities arise mainly where both stimulation sites provide a strong input.

We then asked whether additionally responsive channels were driven by linear sums of weak, subthreshold responses or by nonlinearities. For the additionally responsive TSEP channels, we found 97.8% were classified as linear (with 1.2% supralinear and 1.0% sublinear channels), while for HFA criteria, 90.5% channels were linear (8.0% supralinear, 1.3% sublinear). Additional responsiveness seems therefore driven largely by linear addition of weaker responses, with a small, but meaningful, contribution from supralinearities in HFA additional responsiveness.

### Nonlinear responses vary by region and proximity to stimulation site

Comparing frontal and thalamic-involvement groups, restricted to channels responsive to both stimulation sites (n = 13 and 11 pairs meeting the inclusion threshold), frontal stimulation produced significantly higher rates for both sublinearities (two-sided permutation test; mean difference +10.2%, p = 0.0013, δ = 0.71) and supralinearities (+12.3%, p = 0.013, δ = 0.59; Figure 2h). Thus, multisite stimulation within a cortical structure appears more likely to produce nonlinearities than thalamic or corticothalamic stimulation (nonlinearity rates across regions in Supplementary Figure 2 and Supplementary Table 8).

To assess the effect of proximity to stimulation, we computed the fraction of responsive channels that were nonlinear as a function of minimum distance from either stimulation site (Figure 2i). Sublinearities clustered close to a stimulation site, most within 15 mm, with a sharp drop-off followed by a stable rate. Supralinearities fell off more gradually, with significant supralinear responses as far as 70 mm from either site, suggesting the involvement of long-range network connectivity.

**Figure 2:**
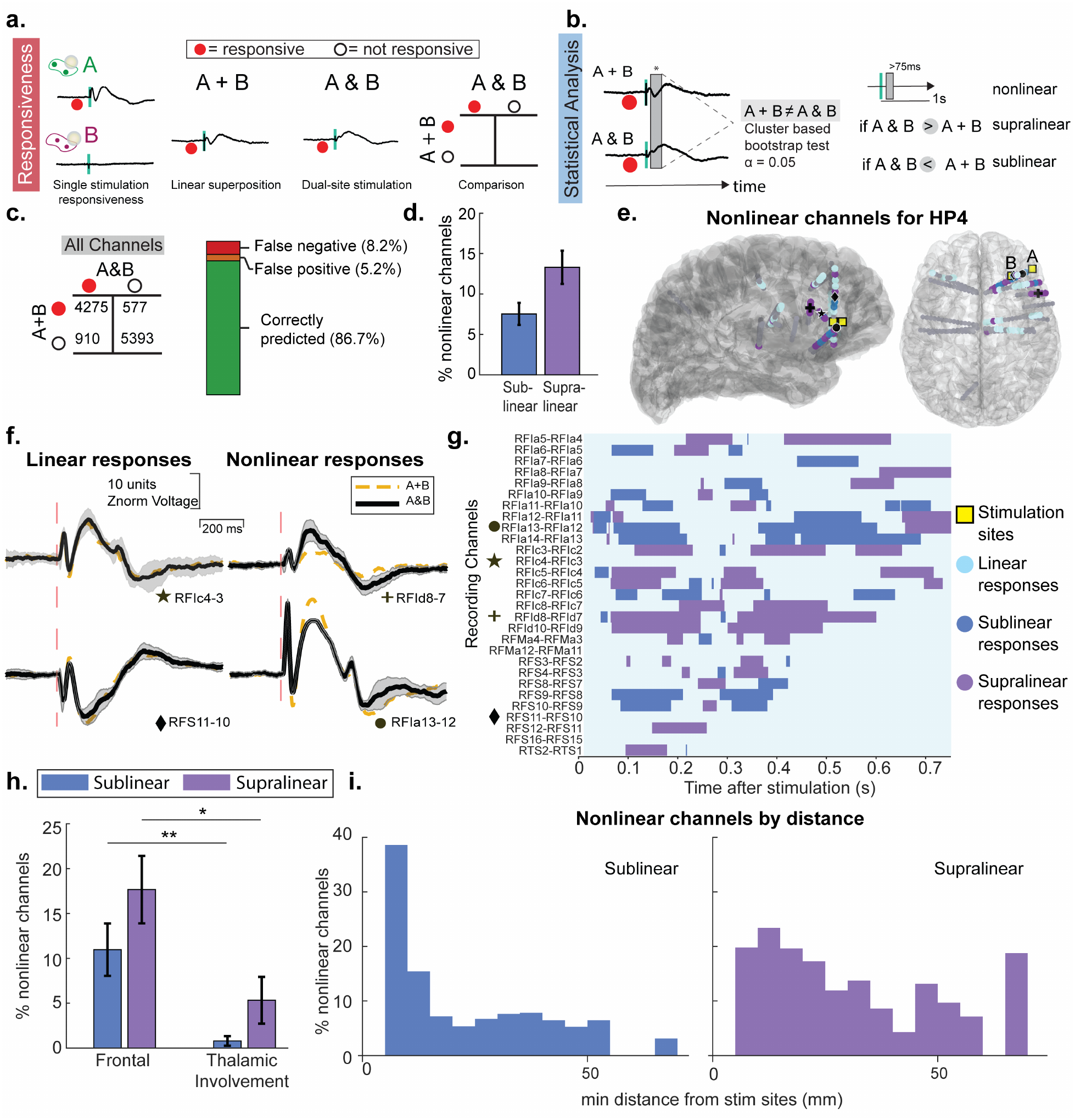
Comparison of dual-site stimulation responses to the linear sum of the two single-site stimulation responses. a, For each stimulation pair, the set of channels responsive to dual-site stimulation is compared with the set predicted to be responsive from the linear sum of the two single-site responses, as a confusion matrix. **b**, Time-resolved comparison of dual-site and linear-sum responses using a cluster-based bootstrap test. Each time point is classified as linear, sublinear, or supralinear. Channels with >75 ms of sublinear (supralinear) activity are classified as sublinear (supralinear). The full rule is given in Methods. **c**, Across all channels at 7 mA, dual-site responsiveness is correctly predicted by the linear sum in 86.7% of cases, with 8.2% false negatives and 5.2% false positives (n = 11,155 channel-pair observations from 87 pairs in 20 participants). **d**, Percentage of channels responsive to both single sites that are classified as sublinear (7.5%) and supralinear (13.3%) at 7 mA. Mean ± s.e.m. across stimulation pairs (n = 63 pairs). e-g, Example stimulation pair from one participant (HP4), with dual-site stimulation through contacts RFIa16-15 (right pars orbitalis) and RFS6-5 (right lateral orbitofrontal cortex). **e**, Spatial distribution of linear and nonlinear channels. Black symbols mark the channels shown in f and gray dots other recording contacts. **f**, Example linear (left) and nonlinear (right) responses. Black traces show the median dual-site response across trials (shading, ±1 s.d.) and mustard dashed traces the sum of the median singlesite responses. **g**, Time course of linear, sublinear and supralinear responses for channels responsive to stimulation at both sites. h, Percentage of channels responsive to both single sites that are classified as sublinear or supralinear, by stimulation location, at 7 mA. Mean ± s.e.m. across stimulation pairs. Two-sided permutation test (n = 13 frontal and 11 thalamic-involvement pairs; *p < 0.05, **p < 0.01). i, Distribution of sublinear (left) and supralinear (right) channels as a function of Euclidean distance from the nearest stimulation site, pooled across all 87 pairs at 7 mA. Bins with fewer than 5 channels responsive to both sites are not shown.

### Nonlinear responses increased with stimulation intensity

Since therapeutic effect is largely pursued by raising stimulation intensity (often at the cost of side effects), we investigated how additional responsiveness and nonlinearity rates varied with intensity. Additional responsiveness was observed at every intensity (Figure 3a). While the TSEP rate showed a small but significant decrease from 4 to 7 mA (paired sign-flip permutation test; mean difference −6.2%, p = 0.023, δ = −0.14, n = 62), the HFA rate did not differ (−4.9%, p = 0.13, δ = −0.04, n = 61). Neither rate differed significantly between 2 and 4 mA (TSEP: −10.6%, p = 0.078, δ = −0.08, n = 20; HFA: −11.0%, p = 0.25, δ = −0.12, n = 13). Additional responsiveness thus persists across stimulation intensities, as it does across brain regions, with only a small decrease at the highest intensity.

Nonlinear responses arise from interactions between the two stimulation responses, and their rate should therefore depend on intensity. Indeed, among channels responsive to both sites, the sublinearities significantly increased from 4 mA to 7 mA (paired sign-flip permutation test; mean difference +6.1%, p = 0.0017, δ = 0.46, n = 30). The supralinearity rate also significantly increased (+6.6%, p = 0.027, δ = 0.40). Two mA was omitted because only two pairs met the minimum-observation threshold (Methods). Higher intensity thus favors nonlinearities.

### Supralinearities were associated with the late TSEP response

Specific trends in timing of nonlinear responses can help explain the underlying mechanisms leading to the observed activity. The early response of the stimulation-evoked potential is believed to reflect local, excitatory activity with the late activity believed to be driven by inhibitory activity^31,39^ and network rebound^40,41^. To characterize the timing of nonlinearities, we generated histograms of the nonlinear time points across channels, averaged over each stimulation pair (Figure 3b). Supralinear time points showed a sharp peak 100-200 ms after stimulation, close to the canonical late peak^30^, whereas sublinear time points were spread more broadly over the response, indicating a process that is less temporally specific.

### Linear superposition predicted TSEP features

A deeper characterization of the physiology underlying responses to dual-site stimulation and how it differs from single-site responses would enable accurate prediction of effects of dual-site stimulation. Neural responses to cortical stimulation can be largely characterized by the area under the curve (AUC), peak amplitude, and timing of its early and late peaks^31,33,42,43^. To test whether these features follow the linear sum across the recorded range, or depart from it beyond a certain linear range of operation, we compared the AUC, amplitude, and time to maximum peak of A&B against those of A+B, pooling channels across participants.

Peak amplitude and AUC were clustered around the linear fit (slope close to 1) for the range of recorded voltages, and across the three stimulation intensities (linear regression at 7 mA; amplitude slope = 0.969, R² = 0.950, nRMSE = 19.9%, Figure 3c; AUC slope = 0.952, R² = 0.907, nRMSE = 31.5%; Figure 3d, all stimulation intensities in Supplementary Table 4). Points furthest from the fit line were mostly nonlinear channels with supralinear above and sublinear below it, and scatter increased with voltage, consistent with the rise in nonlinearity rate at higher amplitudes (Figure 3c). A plateau in these plots, i.e. a level of summed single-site input above which the dual-site response no longer increases, would mark where the response saturates where no further activity can be elicited. However, none was apparent, even at high values of recorded voltages, suggesting linear superposition remained a reasonable prediction across the stimulation and recording amplitudes investigated here (Figure 3c,d).

By contrast, the timing of the maximum peak in the response for ‘A&B’ was not as closely associated with ‘A+B’. While there was clustering of points along the identity line at lower values (corresponding to a dominant early peak), there was a large spread in the plot (linear regression at 7 mA; slope: 0.462, R² = 0.249, nRMSE = 60.8%, Figure 3e). Notably, a distinct cluster of supralinear channels was also seen on the identity line at 100-200 ms. Thus, while AUC and amplitude of stimulation response can be reasonably approximated by ‘A+B’, the timing of the peak response cannot.

### TSEP features and anatomic information were informative predictive features

To identify the most salient features for predicting dual-site responses, we fitted a linear mixed effect model (LME) predicting each dual-site TSEP feature - amplitude, AUC and time to maximum peak - from the features of the summed single-site responses (amplitude, AUC and peak timing), the peak offset (how far, and in which direction, the single-site peaks lie from the summed-response peak), the anatomical group of the stimulation pair (StimLoc) and recording channel (RecordingLoc), and whether the channel lay within 15 mm of a stimulating electrode (isLocal), with participant as the random effect. The TSEP features were z-normalized to index relative importance.

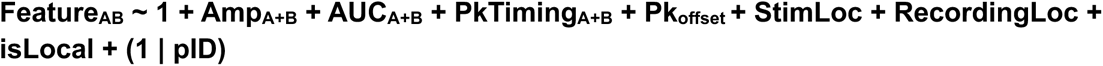

For amplitude, the strongest predictor was the summed single-site amplitude (standardized β = 1.01). Stimulation location, summed AUC, the timing of the summed-response peak and peak offset were also significant. For AUC, the summed AUC dominated (β = 0.78), followed by amplitude, with stimulation location, recording location, and timing also significant. For peak timing, the summed-response peak timing had the largest effect (β = 0.62), followed by the peak offset, with summed amplitude, AUC and stimulation location also significant (full results in Supplementary Table 5).

As per the Akaike Information Criterion (AIC), the complete model was preferred over a one-feature model (**Feature_AB_ ∼ 1 + Feat_A+B_**, equivalent to the linear fits in Figure 3c-d) for the features (Supplementary Table 6), suggesting structural information and recording features can help refine TSEP feature estimates in a given participant. However, under leave-one-participant-out cross-validation the complete model gave marginal to no benefit for amplitude (both models reached R² = 0.95) and AUC (R² = 0.91 vs 0.91), with only modest improvement for time to maximum peak (R² = 0.29 vs 0.24), reflecting the large variability that makes generalization to unseen participants difficult.

**Figure 3:**
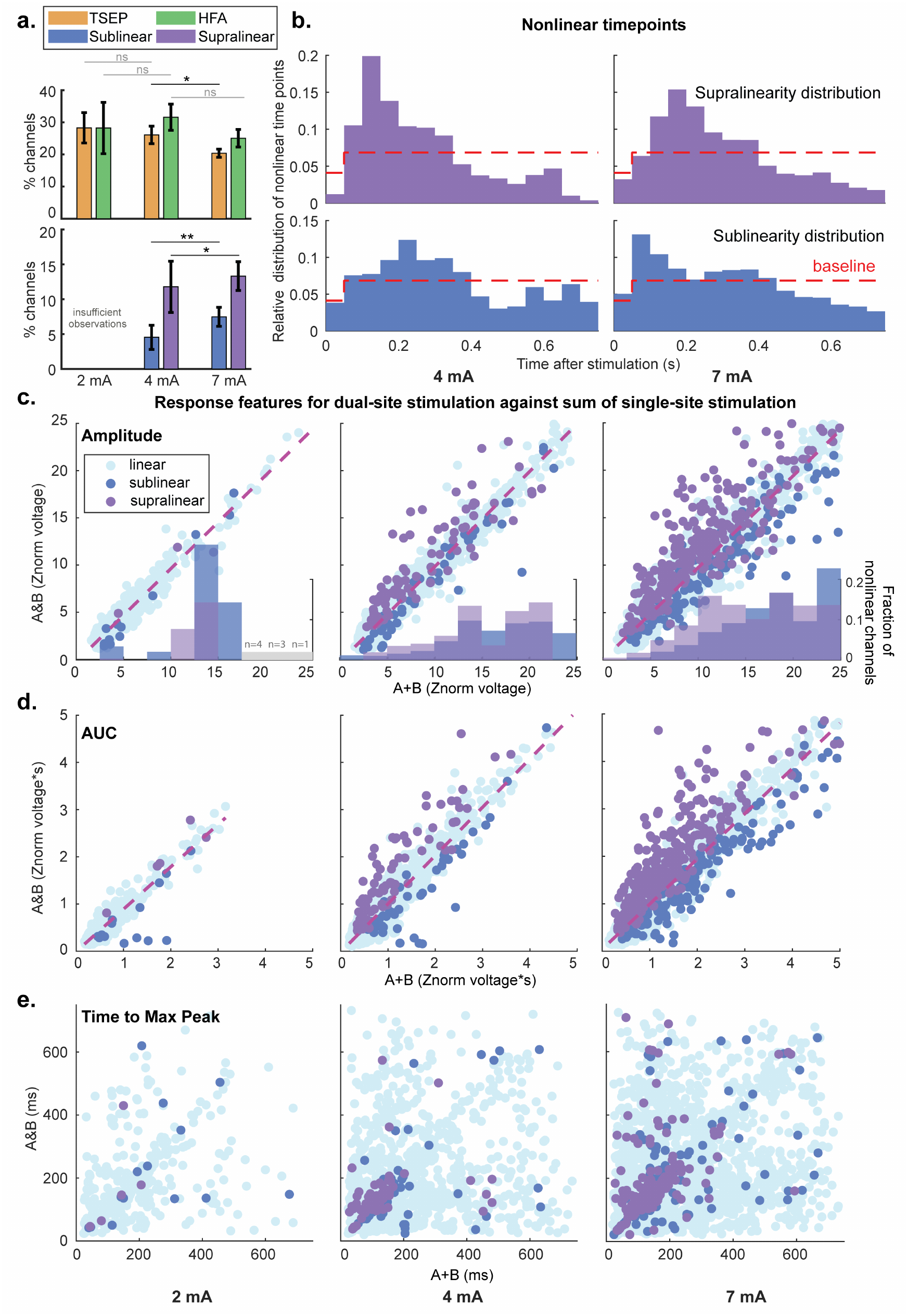
Impact of stimulation intensity on additional responsiveness, nonlinearities and TSEP response features. **a,** Rate of additional responsiveness and nonlinear channels as a function of stimulation intensity. Top: percentage of channels additionally responsive to dual-site stimulation, relative to the union of channels responsive to either single site, by the TSEP (orange; n = 26, 62 and 86 pairs at 2, 4 and 7 mA) and HFA (green; n = 18, 61 and 84 pairs) criteria. Bottom: percentage of sublinear (dark blue) and supralinear (purple) channels among channels responsive to both single sites (n = 30 and 63 pairs at 4 and 7 mA). 2 mA is omitted from nonlinearity because only two pairs met the minimum observation threshold. Mean ± s.e.m. across stimulation pairs. Two-sided paired sign-flip permutation tests on the pairs stimulated at both intensities (4 vs 7 mA: n = 62 TSEP, 61 HFA and 30 nonlinearity pairs; 2 vs 4 mA: n = 20 TSEP and 13 HFA pairs; *p < 0.05, **p < 0.01; ns: not significant). **b,** Weighted histograms of supralinear (top) and sublinear (bottom) time points pooled across channels at 4 mA (left) and 7 mA (right). Distributions are normalized to unit total weight (50 ms bins; red dashed line, uniform baseline of 0.068 per bin (0.041 for the first bin, as testing starts at 20 ms)). 2 mA is omitted owing to too few nonlinear responses. Supralinear time points peak around 100-200 ms after stimulation, whereas sublinear time points are more uniformly distributed across time. **c-e,** Dual-site TSEP features against the corresponding features of the summed single-site responses for channels responsive to at least one single site, pooled across participants, at 2, 4 and 7 mA (left to right; n = 574, 2543 and 4853 channel-pair observations). Light blue, dark blue and purple points denote linear, sublinear and supralinear channels. **c,** Peak amplitude of the dual-site response versus the peak amplitude of the summed single-site responses. Histograms show the fraction of nonlinear channels as a function of summed-response amplitude. Gray bars mark bins with fewer than 5 channels. **d,** Area under the curve (AUC). **e,** Time to maximum peak. Dashed magenta lines in c and d indicate linear fits to all data points. Dual-site response amplitudes and AUC values closely track the linear sum across stimulation intensities, whereas the timing of the maximum peak shows greater scatter around the identity line, partly because the largest peak can switch between early and late components.

### Lower intensity dual-site stimulation matched or exceeded responses to higher intensity single-site stimulation

A key motivation for studying dual-site stimulation is the possibility of engaging a wider network while reducing the stimulation intensity required at any one location. In pairs stimulated at both 4 and 7 mA, we compared response to dual-site at 4 mA with responses to single-site at 7 mA, classifying each channel as *dual-dominant* (the dual-site response exceeded both single-site responses), *single-dominant* (both single-site responses exceeded the dual-site response) or *intermediate* (the dual-site response exceeded one single-site response but not the other, and ties).

Figure 4a shows representative examples of dual-dominant channels. Across channels responsive at both 4 and 7 mA, 18.8% were dual-dominant by AUC, 16.0% single-dominant and 65.2% intermediate (Figure 4c). The proportions by amplitude were similar (19.6%, 10.9%, and 69.5%). Low-intensity multisite stimulation can therefore elicit greater neuronal activity than higher-intensity single-site stimulation in a subset of the stimulation network.

A similar pattern held for dual-site stimulation at 2 mA compared with single-site stimulation at 4 mA: 13.3% of channels were dual-dominant by AUC, 15.6% single-dominant, and 71.1% intermediate (Figure 4b; 13.3%, 12.8%, and 73.9% by amplitude). Together, these data suggest that simultaneous stimulation of multiple regions may help reduce peak stimulation intensity at a given location.

**Figure 4:**
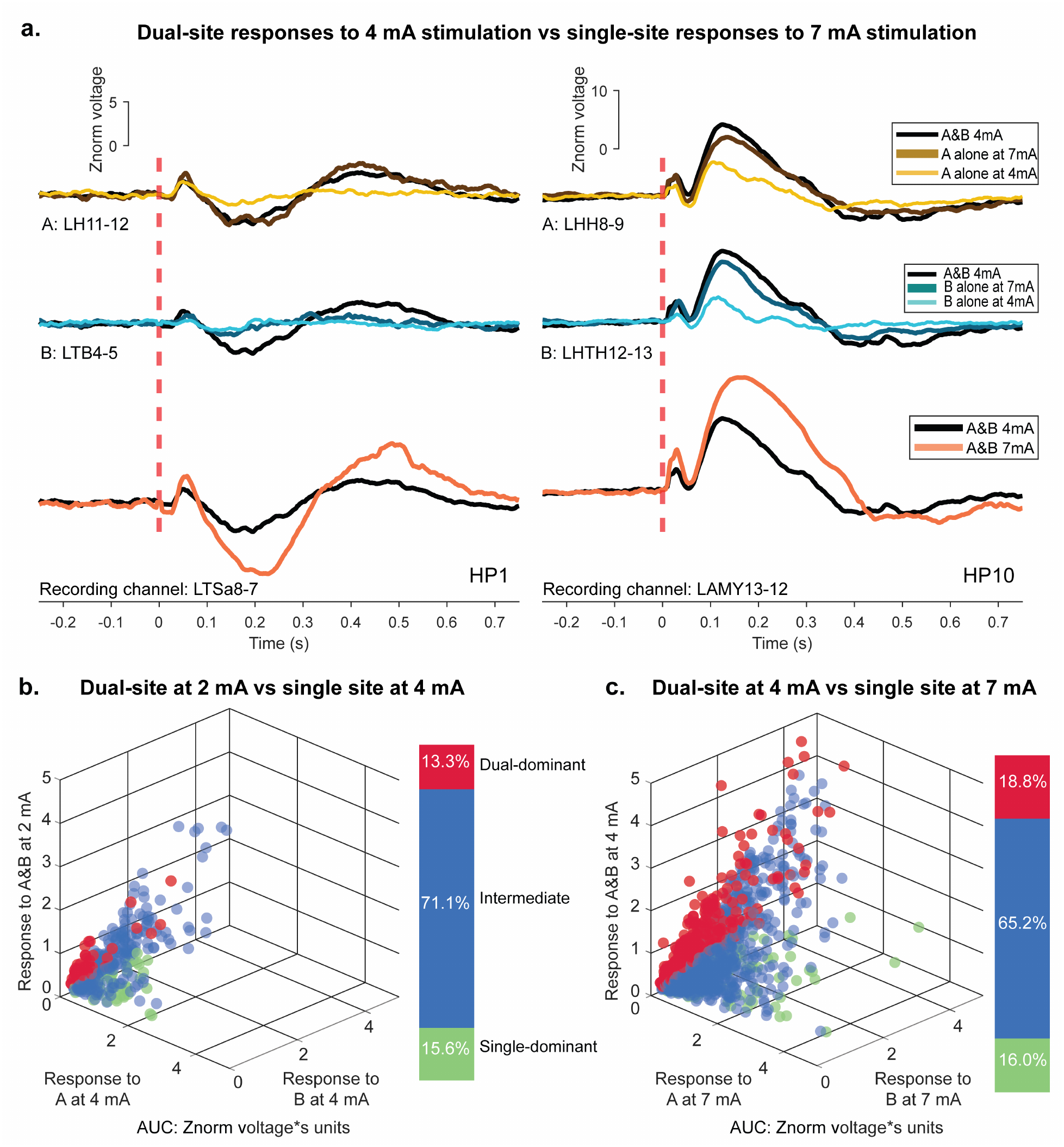
Comparison of effects of dual-site stimulation at a lower intensity with single-site stimulation at a higher intensity. **a**, Example voltage-time responses comparing dual-site stimulation at 4 mA with single-site stimulation at 4 mA and 7 mA for two stimulation pairs. Left, participant HP1 (inferior and middle temporal stimulation, recording channel LTSa8-7) and right, participant HP10 (superior (adjacent white matter) and middle temporal stimulation, recording channel LAMY13-12). Traces show median responses across trials. Light-colored traces show responses to single-site stimulation at 4 mA, and dark-colored traces show responses to single-site stimulation at 7 mA, with black traces showing dual-site stimulation responses at 4 mA. At many recording sites, the response to 4 mA dual-site stimulation matches or exceeds the response to 7 mA single-site stimulation. Bottom, dual-site responses at 4 mA (black) and 7 mA (orange) at the same recording channel. **b,** AUC of the dualsite response at 2 mA against the AUC of the single-site responses at 4 mA at A and B, for channels responsive to at least one single site at both intensities (n = 422 channel-pair observations from 21 pairs in 6 participants). Each point is one recording channel. Red, dual-dominant channels, in which the dual-site response exceeds both singlesite responses. Green, single-dominant channels, in which both single-site responses exceed the dual-site response. Blue points represent the intermediate responses (where the dual-site response exceeds the 4 mA response at one site but not the other, and ties). Stacked bar summarizes the proportions of dual-dominant, intermediate and singledominant responses for AUC. **c,** As in b, dual-site response AUC at 4 mA against the AUC of the single-site responses at 7 mA (n = 2,377 channel-pair observations from 65 pairs in 17 participants).

### Extension to three-site stimulation

To assess whether these results extend to a larger number of concurrent stimulation sites, three-site SPES was delivered to 11 triplets of sites in 7 participants at 4 and 7 mA, with 6 triplets across 5 participants at 2 mA. With two sites, additional responsiveness, driven by the addition of subthreshold responses, was consistent across intensities and locations. If our results extend to multiple sites, the additional responses should be observed at a similar rate. Since sublinearities rose with stimulation intensity, consistent with saturation due to higher activation within a region, a third site should raise the sublinearity rate further, especially at 7 mA. Finally, a third site should drive recorded voltages higher at some locations, testing how far linear superposition extends: a strict suppression threshold would appear as a kink or saturation at the upper end of the three-site amplitude versus A+B+C curve at 7 mA. This saturation should also worsen the performance of the LME model in predicting the three-site stimulation response, with the model assigning a lower weighting to the three-site sum.

### Nonlinearity rates tend to increase with more stimulation sites

Three-site stimulation produced widespread additional responsiveness and nonlinear responses (Figure 5a). To compare their rates with two-site stimulation, we used as the two-site reference the three constituent pairs of each triplet (A&B, A&C and B&C; 32 and 33 pairs against 11 and 11 triplets at 4 and 7 mA (HFA: 31 and 33 pairs); unpaired permutation test; Figure 5b). Additional responsiveness under three-site stimulation was similar to two-site stimulation at every intensity and by both criteria (three-vs two-site). At 4 mA: TSEP 30.9 ± 6.2% vs 29.2 ± 4.5% (p = 0.86, δ = 0.09), HFA 31.1 ± 6.3% vs 31.8 ± 6.5% (p = 0.96, δ = 0.17). At 7 mA: TSEP 23.7 ± 5.0% vs 20.8 ± 2.3% (p = 0.57, δ = 0.05), HFA 34.4 ± 6.3% vs 31.1 ± 4.6% (p = 0.72, δ = 0.17). Not enough triplets met the threshold for HFA (3 of 6) and nonlinearity (0) for 2 mA (all 2-mA values are in Supplementary Table 9). The relatively stable additional responsiveness rate over a larger multisite network suggests that this phenomenon is extendable, and provides a mechanism for accessing broader brain networks with an increase in stimulation sites.

By contrast, among channels responsive to all stimulated sites, nonlinearity rates tended to increase with three-site stimulation. The sublinearity rate showed a non-significant trend tending higher with three sites at 7 mA (14.6 ± 4.2% vs 7.5 ± 1.7%, p = 0.070, δ = 0.33), and similar at 4 mA (7.0 ± 4.3% vs 5.8 ± 2.8%, p = 0.89, δ = 0.09). Supralinearities showed a significant increase at 4 mA (17.9 ± 8.5% vs 5.5 ± 2.0%, p = 0.034, δ = 0.40) but not at 7 mA (19.7 ± 8.2% vs 10.5 ± 2.3%, p = 0.14, δ = 0.13; Figure 5b).

### Linear superposition explains responses to three-site stimulation

Across the three stimulation intensities, we found that three-site peak amplitude remained clustered around the linear fit (slope close to 1) for the range of recorded voltages (linear regression at 7 mA, peak amplitude: slope = 0.93, R² = 0.94, nRMSE = 21.6%; AUC at 7 mA: slope = 0.89, R² = 0.87, nRMSE = 35.1%, Figure 5c). At 7 mA, points at the upper end of recorded activity tended to fall below the fit line, with a sublinearity rate close to 40% at the highest amplitudes. However, no kink or plateau is visible, suggesting there is no clear cutoff where linear superposition fails to explain the responses or where response clearly saturates, though it becomes more likely to be sublinear. The LME analysis for three-site stimulation identified the summed single-site amplitude as the dominant predictor (standardized β = 1.03; Supplementary Table 7). The close agreement of the data with the linear fit (R^2^ = 0.94) and a dominant estimate close to unity suggest linear superposition remains a reasonable hypothesis across the range of recorded voltages.

**Figure 5:**
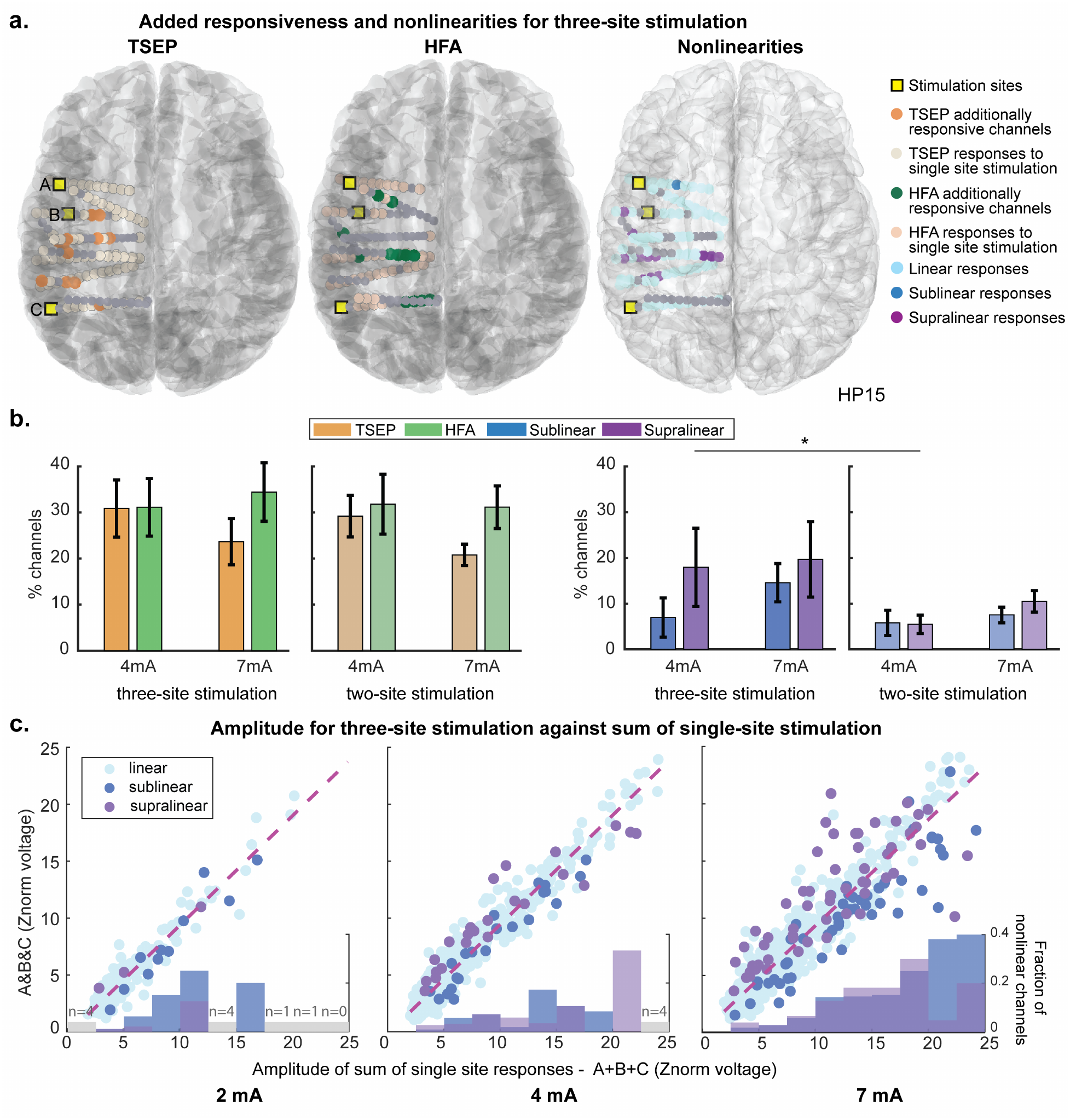
Responses to three-site stimulation. **a**, Example three-site stimulation responses for one participant (HP15) at 7 mA; left to right, TSEP additional responsiveness, HFA additional responsiveness and nonlinearities. Stimulation sites (yellow squares) and recording sites are visualized on a glass brain. Orange and green dots indicate channels that are additionally responsive by the TSEP and HFA criteria, with paler dots showing channels responsive to single-site stimulation. Light blue, dark blue and purple dots indicate linear, sublinear and supralinear channels respectively during three-site stimulation. **b,** Additional responsiveness and nonlinearity rates for threesite and two-site stimulation. Left: percentage of channels additionally responsive to three-site (left) and two-site (right, paler bars) stimulation by the TSEP (orange) and HFA (green) criteria, relative to the union of channels responsive to any single site. Right: percentage of sublinear (dark blue) and supralinear (purple) channels among channels responsive to all stimulated sites, with the same layout. Two-site values are from the three constituent pairs of each triplet (A&B, A&C and B&C). Mean ± s.e.m. across triplets (n = 11 and 11 at 4 and 7 mA; nonlinearity, 5 and 9) and pairs (n = 32 and 33, HFA 31 and 33; nonlinearity, 17 and 28). 2 mA is omitted because too few triplets met the minimum-observation threshold for HFA (3 of 6) and nonlinearity (none). All values are in Supplementary Table 9. Two-sided unpaired permutation tests, three-site vs two-site (*p < 0.05). Unmarked comparisons are not significant. Nonlinearity rates tended to be higher with three sites, with the supralinearity rate significantly higher at 4 mA. c, Three-site TSEP peak amplitude against the peak amplitude of the summed singlesite responses (A + B + C), pooled across participants, at 2, 4 and 7 mA (left to right; n = 185, 570 and 839 channeltriplet observations). Each point is one channel, colored as in a. The dashed magenta line is the linear fit to all data points. Histograms indicate the fraction of nonlinear channels as a function of the summed-response amplitude. Gray bars mark bins with fewer than 5 channels.

## Discussion

In this study, we characterized the physiology underlying responses to synchronous stimulation delivered to multiple locations in the human brain. Across brain regions and stimulation intensities, dual-site response magnitudes were largely explained by linear superposition, while additional responsiveness was obtained. Additional responsiveness was largely consistent with the linear addition of weaker, sub-threshold responses, with distinct nonlinear responses observed mainly at sites effectively connected to both stimulation locations. Within the nonlinear effects, we identified two patterns of nonlinear responses, sublinearities, likely related to saturation and local activity, and supralinearities, arising from late responses and probing inhibitory networks. Together, these findings help explain the relative benefit a second stimulation site contributes to the network engaged. Additional responsiveness can arise through summation, while nonlinear interactions shape the combined response at particular locations and times. We found these principles extended to three-site stimulation, with a tendency toward greater nonlinearity rates.

### Dual-site stimulation can broaden network engagement

The observed additional responsiveness shows that synchronized dual-site stimulation can access a larger effective network than when stimulating one site at a time, as reflected in both recorded TSEPs and rise in local neuronal firing (captured by HFA). This additional responsiveness is largely consistent across stimulation intensities, locations, and number of sites stimulated (Figures 1f, 3a, 5b). Additional responsiveness appears largely driven by a linear addition of weaker, sub-threshold responses: additionally responsive sites are seen distal to the stimulation channels rather than proximal (Figure 1g), where single-site responses, whose amplitude falls with distance^38^, are less likely to cross the threshold for responsiveness, but their sum does. There is a small contribution to this effect from supralinearities, but the bulk of additionally responsive channels are classified as linear in our analysis, which suggests that these occur at locations weakly connected to both stimulation locations. The similar rates of additional responsiveness across intensities suggest that a 20-25% larger brain network can be engaged under multisite stimulation likely due to synchronization of the stimuli.

### Nonlinearities likely reflect strong network interactions in the cortex

Distinct nonlinear interactions in response to dual-site stimulation were observed across participants and stimulation regions, primarily at sites effectively connected to both locations (Figure 2d). Nonlinear responses likely reflected strong interactions elicited by the two (or more) stimulations, and were also influenced by stimulation intensity and recorded amplitude, with greater likelihood of nonlinear responses for higher intensities or amplitudes (Figure 3). Cortical stimulation within the frontal lobe was more likely to elicit nonlinear responses than thalamic or thalamocortical stimulation, though more observations are needed to address these effects across brain regions. The predominance of linear responses at commonly responsive sites is also consistent with the additive interactions reported for clinical high-frequency multi-target stimulation^44^.

These nonlinear interactions seemed to arise from two distinct phenomena, with sublinearities likely linked to local saturation and supralinearities to network inhibitory interactions. Indeed, sublinear responses were mostly proximal to a stimulating electrode, were less specific in timing (Figures 2i, 3b), and their rate increased with stimulation intensity (Figure 3a). Thus, sublinearities may arise from a saturation effect; at a given location, the local neuronal populations may be excited to the point of saturation by one stimulus such that a second stimulus is unable to drive further activity or driven to a refractory state by the first stimulus where the second stimulus is unable to cause further responsiveness. On the other hand, supralinear responses appear to be driven by a more mechanistically specific phenomenon linked to the canonical late stimulation-evoked response. Supralinearities occur most often 100-200 ms after stimulation (Figure 3b), i.e. responses most frequently exceed the sum during this window. Interestingly, we also observed a large cluster of supralinear responses whose peak timing is 100-200 ms after stimulation (Figure 3e), i.e. the peak of the response for supralinear channels lines up with the period that the response exceeds the sum. This suggests supralinear channels appear to have a dominant late peak, which is being strengthened or sustained with dual-site stimulation. Supralinearities are observed at a significant rate distant from both stimulation electrodes, as far as 70 mm (Figure 2i). Since a dominant late peak is typically associated with distant or network activity^31,33,37,38,41^, this suggests a possible role of network activity in generating supralinear responses.

Deeper investigation is needed to verify the proposed link to physiology underlying supra and sublinearities. One option would be to conduct single unit recordings for multiple stimulations to relate neuronal firing and cell types with the types of observed nonlinearity (since the late response is often associated with inhibitory activity^45^). Alternatively, given the specific temporal peaks of supralinearities, varying the delay between delivery of the two pulses may provide a mechanism for selectively inducing these effects, similar to paired-pulse paradigms^46^.

### Dual-site stimulation responses can be accurately predicted from single-site responses

Linear superposition predicted which channels would respond under dual-site stimulation (with 86.7% accuracy) and the amplitude and AUC of the response. This held across intensities and for three-site stimulation (Figures 2c, 3c,d, 5c; Supplementary Tables 5, 7). The success of the linear superposition also indicates what a second stimulation site would contribute in clinical multi-target strategies: where two sites were effectively connected to common regions, synchronized stimulation would engage a broader network and elicit interactions at commonly responsive sites, whereas sites without shared connectivity would behave somewhat independently, consistent with a randomized comparison in obsessive compulsive disorders in which combined stimulation of two targets in distinct prefrontal networks added little over either alone^47^.

Inherent to DES studies, response variability was large across stimulation sites, recording sites, and individuals. The LME analysis investigated whether access to additional features, information about stimulation and recording regions, and accounting for participant variability would improve performance in predicting a given TSEP feature (amplitude, AUC, time to max peak). The complete model was preferred to the simple model, with other TSEP features and stimulation region consistently important. However, when evaluated on held-out participants, the complete model provided marginal benefit over linear superposition alone for amplitude and AUC. A modest gain was obtained for peak timing, which remained poorly predicted by either model. Models that generalize across individuals will likely require additional complexity or features, such as structural and functional connectivity, to help capture intrinsic network properties. Future studies, with larger cohort sizes, may help link the variability of multisite responses to factors known to shape single-site stimulation responses^37,38,48–51^.

### Multisite stimulation can help reduce peak stimulation intensities

We showed that dual-site stimulation at a lower intensity can match or exceed single-site stimulation at a higher intensity. Similar TSEP responses were produced, with features exceeding those at a single-site with higher stimulation current. However, a basic stimulation threshold must be crossed to see particular effects or interactions, as underscored by the lack of sufficient channels responsive to single-site stimulation of both sites at 2 mA. A further study involving balanced conditions could help determine whether multisite stimulation reduces the net current required or primarily redistributes it, and to connect electrophysiology spread to clinical effects. Nevertheless, these data do show multisite stimulation usefulness in reducing peak stimulation intensities, which could help address side effects associated with large stimulation current intensities^52,53^ by engaging multiple nodes at lower currents.

Additional limitations should be noted. These phenomena were studied in brains of patients with epilepsy of heterogeneous pathologies. Inherent to any sEEG investigation, epileptogenicity cannot be ruled out as a contributing factor for the variation observed. To mitigate this, we excluded channels considered epileptic by either an automated algorithm^54^ or classified as being in the seizure onset zone. Our analysis is also less sensitive to nonlinearities in the early response, since the first 50 ms combines the physiological early peak with blanking and any residual stimulation artifact, so that small deviations from the linear sum during this period of high variance are unlikely to reach significance in the cluster-based analysis; a larger number of trials would be needed to resolve early effects. Further, due to the low number of triplets stimulated for the three-site stimulation studies, any statistical observations related to three-site stimulation should be treated as tentative. Finally, we characterized responses to synchronous single pulses, and further work is needed to establish how these principles extend to the stimulation trains used therapeutically and to behavioral or clinical outcomes.

More broadly, our results help provide a concrete physiological understanding of the interactions resulting from multisite stimulation in humans. At the scale of the brain’s effective networks, simultaneous inputs to two nodes combined largely linearly, with departures that marked possible inhibitory network interactions (supralinear, late, distant) and saturation (sublinear, less temporally specific, local). As clinical stimulation moves towards targeting multiple nodes and engaging disease networks, this work represents a foundational step towards principled, network-guided neurostimulation.

## Acknowledgement

We would like to thank the patients who participated in this research for their time and help. This research was supported by the National Institutes of Health (NIH) grants 1RF1NS132784, 5R01AG090302, and 1S10OD034414. The work was also supported by the MIT School of Engineering MathWorks Fellowship.

## Code and Data Availability

All code used for data analyses in this study will be made available on the MGH Center for Neurotechnology and NeuroRecovery GitHub (https://github.com/Center-For-Neurotechnology) with the publication of this manuscript. The fully de-identified raw data associated with this study will be made available on the DABI repository in the iEEG-BIDS^55^ format with the publication of this manuscript.

## Methods

### Experimental Model and Participant Details

Twenty patient participants with semi-chronic depth electrodes participated in the study (10 female; age 21–63, mean 37.0 years; 3–6 pairs per participant). All participants received stimulation at 7 mA (87 pairs); 17 participants (66 pairs) also received stimulation at 4 mA and 7 participants (27 pairs) at 2 mA. Three-site stimulation was conducted in 7 of these participants (11 triplets at 4 mA, 11 at 7 mA; 6 triplets in 5 participants at 2 mA). The stimulation was conducted in the epilepsy monitoring unit (EMU). The participants were aware they were receiving stimulation, but blinded to timing, location and intensity of the stimulation.

Electrodes were implanted exclusively for clinical reasons at the Massachusetts General Hospital (MGH). Depth electrodes (PMT, Chanhassen, MN, USA, or DIXI) were stereotactically placed in locations deemed necessary for seizure localization by a multidisciplinary clinical team independent of this research. Following implant, the preoperative T1-weighted MRI was aligned with a postoperative CT or MRI using volumetric image co-registration procedures and FreeSurfer scripts^56^. Electrode coordinates were manually determined from the CT or MRI in the participants’ native space. Each contact was assigned an anatomical region from a standardized cortical map, using an electrode labeling algorithm. The image processing pipeline is described in detail in Soper et al.^57^.

All patients voluntarily participated after fully informed consent as monitored by the Institutional Review Board (IRB) of Mass General Brigham which oversees both Massachusetts General Hospital (MGH) and Brigham and Women’s Hospital (BWH). Participants were informed that participation in the stimulation experiment would not alter their clinical treatment and that they could withdraw at any time without jeopardizing their clinical care.

### Neural stimulation and recordings

Single pulse electrical intracranial stimulation was delivered with a CereStim R96 stimulator (Blackrock Microsystems, Salt Lake City, UT) while simultaneously acquiring intracranial EEG (iEEG) with either Blackrock Neural Signal Processing system at a 2 kHz sampling rate (Blackrock Microsystems, Salt Lake City, UT, USA), or a Neuralynx Atlas system at 2 kHz sampling rate. Depth recordings were acquired with reference to an EEG electrode placed on the scalp (either O2 or O1). Individual charge-balanced biphasic stimulation pulses were 233 µs duration, 90 µs long each phase with 53 µs interphase interval. The current intensity was either 2 mA, 4 mA or 7 mA, the interstimulation interval was 4-5 s with ±0.25 s jitter. Current injection and return paths used neighboring contacts in a bipolar configuration.

For stimulation, a stimulation group of three distinct sites was chosen. Stimulation was pseudo-randomly delivered to the three stimulation sites and the three dual-site pairs, controlled via a custom Cerestim API (https://github.com/Center-For-Neurotechnology/CereLAB) via MATLAB at one intensity for 20 trials per stimulation site. This results in delivery of stimulation in 6 distinct conditions (site ‘A’, site ‘B’, site ‘C’, site ‘A’ and ‘B’ together, site ‘A’ and ‘C’ together, site ‘B’ and ‘C’ together) for a stimulation group. This was then either repeated for another intensity, or time permitting, for another group of three sites. Each participant received stimulation for at least one group of three sites, while some participants received stimulation at two groups (no participant received stimulation to more than two groups at a given intensity). In the participants who received three-site stimulation, all three sites were also stimulated together (A&B&C) as a seventh block at the same intensities and trial count. A trained epileptologist/electroencephalographer examined continuous recordings for epileptiform activity, and participants were asked if they experienced any sensations. No participants reported any discomfort or sensations with the stimulation delivered, and no seizures occurred during or shortly after the stimulation.

### Choice of stimulation sites

The stimulation sites were chosen to maximize the number of recording channels which were responsive to more than one stimulation site at a time. This was to ensure we could study the interactions at a given region due to stimulation of two sites, since our initial hypothesis suggested that recording channels which were responsive to stimulation of only one of the sites would be relatively unaffected by simultaneous stimulation of a second stimulation location which was not effectively connected to the recording channel.

To achieve this, we studied the response to single-site SPES of a large number (20-60) of cortical and subcortical sites (similar to protocol described in^37,38^). Using the TSEP responsiveness criteria, we identified the set of responsive channels for each site, and ranked candidate stimulation pairs based on descending order of number of common responsive channels. Potential pairs were excluded if they elicited epileptiform activity in response to stimulation over a large number of channels (manually), or if both sites were on the same shaft (to remove the confound of intra-shaft effects).

It was not always possible to run the single-site stimulation survey experiment. In those cases, we chose sites based on the location which would have stronger responses and broad connectivity, and therefore a larger possibility for having common responsive channels, focusing on sites in the gray-white boundary^37^.

### Data Preprocessing

To account for epileptiform activity, channels with pathological interictal epileptiform discharges (IEDs) were excluded from analysis using an automated IED detection algorithm specified by Janca et al^54^, as were channels highlighted to be in the seizure onset zone as per clinical reports. Further, channels with excessive artifact, railing or noise, determined by visual inspection, were also removed from analysis.

Bipolar re-referenced data were used for analysis. Stimulation artifacts were removed by blanking the signal from 5 ms before to 15 ms after each stimulus onset, with the blanked window reconstructed using cubic spline interpolation anchored on 5 ms of clean data on either side of the blanked period, adapted from David et al.^58^. The line noise was removed using a 60 Hz (and 120 Hz) notch filter. The data were low pass filtered to 250 Hz and downsampled to 500 Hz since that captured the frequencies of interest. Activity in each channel was z-normalized with respect to a 10 second baseline period before the start of stimulation per channel. Stimulation in each condition was conducted for 20 trials, and an epoch of 1.5 s before and after stimulation onset was used for analysis, and the signal was demeaned per trial. This approach ensured that the signals in the single-site and dual-site conditions were scaled by the same baseline so could be compared, but demeaned per trial to account for any drift. All analysis was conducted in MATLAB 2021a. The authors used LLMs (Claude and ChatGPT) to assist with refactoring of the analysis code and for proofreading of the manuscript. All code and text were verified by the authors, who take full responsibility for the content.

### Quantification and Statistical Analysis

#### TSEP detection

Responsive channels were identified using the algorithm of Zelmann et al.^48^ with thresholds relaxed to increase sensitivity. For each bipolar channel, the trial-wise median of the baseline-normalized response was computed, and positive and negative peaks were detected post-stimulation (minimum prominence 1, minimum width 12.5 ms, minimum inter-peak interval 25 ms). Peak, peak-to-peak and peak-to-peak-to-peak amplitudes and their corresponding areas (prominence × half-peak width) were extracted. A null distribution for each feature was built from 100 randomly placed pre-stimulation segments of equal duration, and the 95th percentile defined the channel-specific baseline thresholds. A channel was classified as responsive if, for any feature family, its amplitude or area exceeded 2 × the corresponding baseline threshold while also exceeding absolute minima of z = 1.96 (corresponding p < 0.05) and AUC = 100 z·ms, with the first peak occurring within 500 ms of stimulation. Channels with fewer than 5 artifact-free trials were excluded. The same detector and criteria were applied to the summed single-site responses (A+B) to obtain the predicted responsive set used in Figure 2a,c.

The key features of the TSEP used in our investigations were the amplitude of the response, area under the curve and timing of maximum peak. The amplitude was quantified using the peak-to-peak voltage, computed as the difference between the minimum and maximum z-normalized voltage recorded from 20 ms to 750 ms post stimulation. The area under the curve computed the integral of the absolute value of the response for 20 ms to 750 ms post stimulation, while the time to maximum peak refers to the time in ms corresponding to the largest peak (absolute deflection) in this window.

#### Spectral Analysis

A multitaper spectrogram of the response was computed for each channel using the chronux^59^ mtspecgramc function (100 ms moving window, 10 ms step, time-bandwidth product of 2 with 3 tapers), and the activity from 70-150 Hz, removing a 10 Hz band around 120 Hz, expressed relative to the pre-stimulus baseline power, was used to assess whether there was a rise in local neuronal firing in response to stimulation. A channel was considered HFA-responsive when power in the HF band crossed a threshold of 2.5 times baseline and had an AUC above 1 exceeding 0.15. For reference, a peak of 2.5 for at least 100 ms meets this AUC. Having an AUC criterion includes both cases where there may be a sharp rise for a shorter time period, or a smaller rise for a longer time period, while having a minimum threshold of activity ensures robustness to small baseline shifts.

#### Nonlinearity Classification

For each recording channel we tested whether the response to simultaneous dual-site stimulation (AB) differed from the linear sum of the two single-site responses (A + B). We did this by computing the difference between the means at each time point, D(t) = mean(AB) – {mean(A) + mean(B)}, and expressed it as a Welch t-statistic^60^. Significance was assessed with a cluster-based test^61^: adjacent time points exceeding a pointwise threshold (two-sided p < 0.05) were grouped into clusters, and each cluster was summarized by its mass, the sum of its t-values. The null distribution of cluster mass was obtained by bootstrap resampling.

Each condition’s response was de-meaned, so that the three conditions were exactly additive while retaining their own variability, and whole trials were then resampled with replacement within each condition^62^. In each of 10,000 resamples, the largest cluster mass was recorded, and a cluster in the data was considered significant if fewer than 5% of resamples produced a larger one, controlling false positives across all tested time points of a channel (20–750 ms after stimulation). Within significant clusters, each time point was labelled supralinear when AB and A + B had the same sign and AB was larger in magnitude, and sublinear when it was smaller. Time points where their signs differed were excluded. A channel was classified as supralinear (sublinear) if its supralinear (sublinear) time points totalled more than 75 ms. For three-site stimulation, the difference was mean(ABC) − mean(A) − mean(B) − mean(C).

Across TSEP, HFA and nonlinearity criteria, rates are reported as mean ± s.e.m. across stimulation pairs (or triplets) unless stated otherwise.

#### Linear Mixed Effect Models

We used Linear Mixed Effect (LME) models to identify the key features involved in influencing the response to dual-site stimulation, analogous to approaches described in Kundu et al. and Zelmann et al.^48,63^ using participant identity as the key random effect. We tested the complete model:

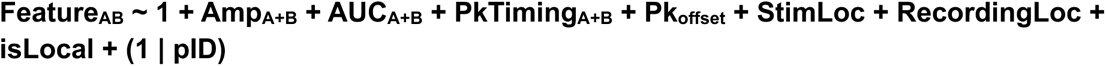

Where the subscript AB corresponds to the dual-site stimulation response, and A+B to the features generated from the sum of the medians of the single-site responses. The amplitude, AUC and peak-timing features of the A, B, A&B and A+B responses, and peak offset, were z-normalized before fitting so that standardized coefficients are comparable across features. Channels not responsive to stimulation were omitted from the LME analysis.

Peak offset quantifies how the timing of the single site peaks differs from the timing of the peak for the summed response, weighted by their amplitudes. It is 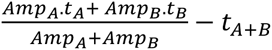, where t is the time to max peak, and Amp is the peak amplitude of the response. Since timing doesn’t appear to cleanly follow the sum, this term aims to capture the misalignment between the timing of the single site peaks and the summed response, and explores whether that helps the linear model improve its performance on predicting timing of the maximum peak for the dual-site response.

StimLoc is a categorical feature derived based on the location of the stimulation sites; The core groups of interest used were: 1) frontal (both sites in the frontal lobe in the same hemisphere); 2) temporolimbic (both sites in the temporal lobe or limbic areas in the same hemisphere); 3) white matter (both sites in white matter); 4) homologous (both sites in homologous locations in the two hemispheres); 5) thalamic involvement (one or both of the stimulation location is in the thalamus); 6) other.

RecordingLoc is a categorical feature derived on the broad anatomic structure the recording channel is present in^48^. The following regions of interest were defined and compared: i) frontal, ii) temporal, iii) Central Sulcus, iv) Parietal (including parietal and occipital), v) Limbic, vi) Insula, vii) Subcortical (Thalamic, Caudate, Accumbens, and Putamen), viii) white matter, ix) unknown.

Note the temporolimbic stimulation group is broken into its constituent groups for recording locations, since there are sufficiently many observations per group (frontal 1390, temporal 508, central sulcus 145, parietal-occipital 65, limbic 185, insula 128, subcortical 326, white matter 1815, unknown 289). Because several stimulation pairs were tested in each participant, each recording channel contributes one observation per stimulation pair. Pooled counts are therefore channel-pair observations (2561 unique bipolar channels across the 20 participants).

isLocal is a binary flag expressing whether the recording location is within 15 mm of a stimulating electrode, since stimulation evoked response amplitudes decline with distance from the stimulation site with an inflection near 15 mm^37^ which separates near-field local responses from propagated distant activity^38^.

#### Permutation tests

Group comparisons were assessed using two-sided permutation tests (10000 permutations), with the test statistic being the difference in group means, and a null distribution was built by pooling all observations and randomly reassigning them to two groups matching the original sizes. For paired comparisons (the same stimulation pairs at two intensities), we used a sign-flip permutation test on the within-pair differences, in which the sign of each difference was independently flipped with equal probability at each permutation to build the null distribution. In both cases the two-sided p-value was the proportion of permutations whose absolute statistic equaled or exceeded the absolute observed value, and p = 0.05 was used for assessing significance. For each comparison we also report Cliff’s δ as a distribution-free effect size. Positive mean differences and δ indicate higher values in the first-named group or, for intensity comparisons, at the higher intensity.

A stimulation pair (or triplet) contributed to a rate only if at least five channels met the criterion defining that rate’s denominator: for additional responsiveness, at least five channels responsive to either single site by the TSEP or HFA criterion, respectively, while for nonlinearity rates, at least five channels responsive to both (all) stimulated sites. Rates are reported as mean ± s.e.m. across the contributing pairs or triplets. Nonlinearity rates at 2 mA are not reported because only two pairs and no triplet met this threshold.

