## Supplementary Information for "Electrophysiology of multisite direct electrical stimulation in the human brain"

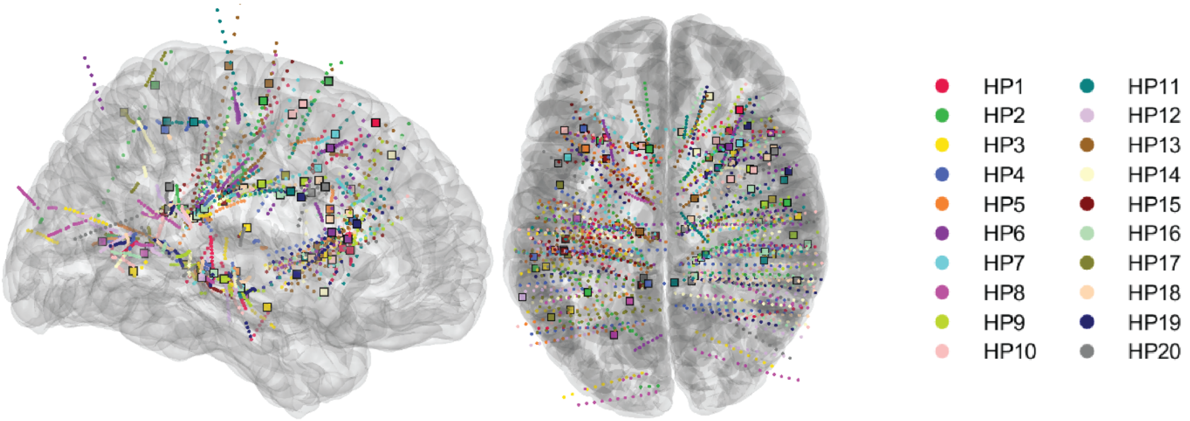


**Supplementary Figure 1:** Stimulation and recording locations pooled across participants on an MNI brain.

| Supplementary Table 1: Stimulation locations across participants | | | | | | | | | |
|---|---|---|---|---|---|---|---|---|---|
| Participant ID | Stimulation Pair | Stimulation Amplitude (mA) | Electrode Pair 1 | Electrode Pair 2 | Electrode Pair 3 | Electrode Region 1 | Electrode Region 2 | Electrode Region 3 | Three-site Stimulation conducted? |
| HP1 | 1 | 4, 7 | LH11-LH12 | LTB04-LTB05 | RFI09-RFI10 | L-temporal | L-temporal | R-frontal | No |
| | 2 | 7 | RFI09-RFI10 | RTP08-RTP09 | RTSa7-RTSa8 | R-frontal | R-temporal | R-temporal | No |
| HP2 | 1 | 4, 7 | LFIa13-LFIa14 | LPIb3-LPIb4 | LTSb7-LTSb8 | L-frontal | L-limbic | L-temporal | No |
| HP3 | 1 | 4, 7 | LAMY03-LAMY04 | LFIb10-LFIb11 | LTSb5-LTSb6 | L-limbic | L-WM | L-temporal | No |
| HP4 | 1 | 4, 7 | RFIa15-RFIa16 | RFMa06-RFMa07 | RFS05-RFS06 | R-frontal | R-frontal | R-frontal | No |
| HP5 | 1 | 4, 7 | LFI07-LFI08 | LFMb15-LFMb16 | RFMb7-RFMb8 | L-frontal | L-frontal | R-frontal | No |
| HP6 | 1 | 4, 7 | LAMY09-LAMY10 | LHT02-LHT03 | LTB05-LTB06 | L-WM | L-limbic | L-occipital | No |
| HP7 | 1 | 4, 7 | RFIcA12-RFIcA13 | RFMaC13-RFMaC14 | RHH10-RHH11 | R-frontal | R-WM | R-temporal | No |
| HP8 | 1 | 7 | LFM02-LFM03 | RFMa08-RFMa09 | RFMb08-RFMb09 | L-frontal | R-WM | R-WM | No |
| | 2 | 7 | LFM13-LFM14 | RFMa08-RFMa09 | RFMb11-RFMb12 | L-frontal | R-WM | R-frontal | No |
| HP9 | 1 | 2, 7 | RFMa5-RFMa6 | RFMbA15-RFMbA16 | RPI1-RPI2 | R-frontal | R-frontal | R-limbic | No |
| | 2 | 2, 7 | RFMbA15-RFMbA16 | RFOCM3-RFOCM4 | RPI11-RPI12 | R-frontal | R-thalamus | R-WM | No |
| HP10 | 1 | 2, 4, 7 | LHH8-LHH9 | LHTH12-LHTH13 | RHH8-RHH9 | L-WM | L-temporal | R-WM | No |
| HP11 | 1 | 4, 7 | LFIb09-LFIb10 | LFSd06-LFSd07 | RFSb05-RFSb06 | L-frontal | L-WM | R-frontal | No |
| | 2 | 4, 7 | LFIb09-LFIb10 | LFIcA01-LFIcA02 | RFIa06-RFIa07 | L-frontal | L-thalamus | R-WM | No |
| HP12 | 1 | 4, 7 | RFMa1-RFMa2 | RFMcCM2-RFMcCM3 | RFSa8-RFSa9 | R-frontal | R-thalamus | R-frontal | No |
| HP13 | 1 | 7 | LFIa07-LFIa08 | LFMA12-LFMA13 | LPIP11-LPIP12 | L-frontal | L-frontal | L-WM | No |
| | 2 | 7 | LFIa07-LFIa08 | LFMA01-LFMA02 | LPIP01-LPIP02 | L-frontal | L-thalamus | L-thalamus | No |
| HP14 | 1 | 4, 7 | RFIcA10-RFIcA11 | RTSaC3-RTSaC4 | RTSbP2-RTSbP3 | R-frontal | R-thalamus | R-thalamus | Yes |
| | 2 | 4, 7 | RAMY7-RAMY8 | RHH13-RHH14 | RTSaC10-RTSaC11 | R-WM | R-temporal | R-temporal | Yes |
| HP15 | 1 | 2, 4, 7 | LFIANT13-LFIANT14 | LHH7-LHH8 | LPI13-LPI14 | L-frontal | L-WM | L-parietal | Yes |
| HP16 | 1 | 2, 4, 7 | RFIa6-RFIa7 | RFIbANT3-RFIbANT4 | RFIc13-RFIc14 | R-frontal | R-thalamus | R-frontal | Yes |
| | 2 | 4, 7 | RFIa6-RFIa7 | RFIbANT15-RFIbANT16 | RFIc2-RFIc3 | R-frontal | R-frontal | R-limbic | Yes |
| HP17 | 1 | 2, 4, 7 | LFSb07-LFSb08 | LPSO12-LPSO13 | LTSPUL13-LTSPUL14 | L-frontal | L-parietal | L-temporal | Yes |
| | 2 | 2, 4, 7 | LFIa09-LFIa10 | LFSb07-LFSb08 | RFMaA15-RFMaA16 | L-frontal | L-frontal | R-frontal | Yes |
| HP18 | 1 | 2, 4, 7 | RFIa11-RFIa12 | RFIb3-RFIb4 | RFIcANT15-RFIcANT16 | R-frontal | R-frontal | R-frontal | Yes |
| HP19 | 1 | 4, 7 | LFIA13-LFIA14 | RFIbA13-RFIbA14 | RTSbP13-RTSbP14 | L-frontal | R-frontal | R-WM | Yes |
| | 2 | 4, 7 | RFIbA1-RFIbA2 | RTSbP1-RTSbP2 | LTSP01-LTSP02 | R-thalamus | R-thalamus | L-thalamus | Yes |
| HP20 | 1 | 2, 4, 7 | LifOF8-LifOF9 | RifOF7-RifOF8 | RmfCD11-RmfCD12 | L-frontal | R-frontal | R-frontal | Yes |

**Supplementary Table 1:** Stimulation locations across participants

For each participant, stimulation was delivered across one or more bipolar electrode pairs (Pair). Amplitudes indicate the current levels (mA) tested for each configuration. Each stimulation site is listed as a bipolar contact pair (e.g., LH11-LH12), where the first contact is the reference used for anatomical localization and the second is the adjacent higher-numbered contact on the same lead. Regions 1-3 give the anatomical assignment for Electrode Pairs 1-3, respectively. Hemisphere is indicated by the prefix L (left) or R (right).

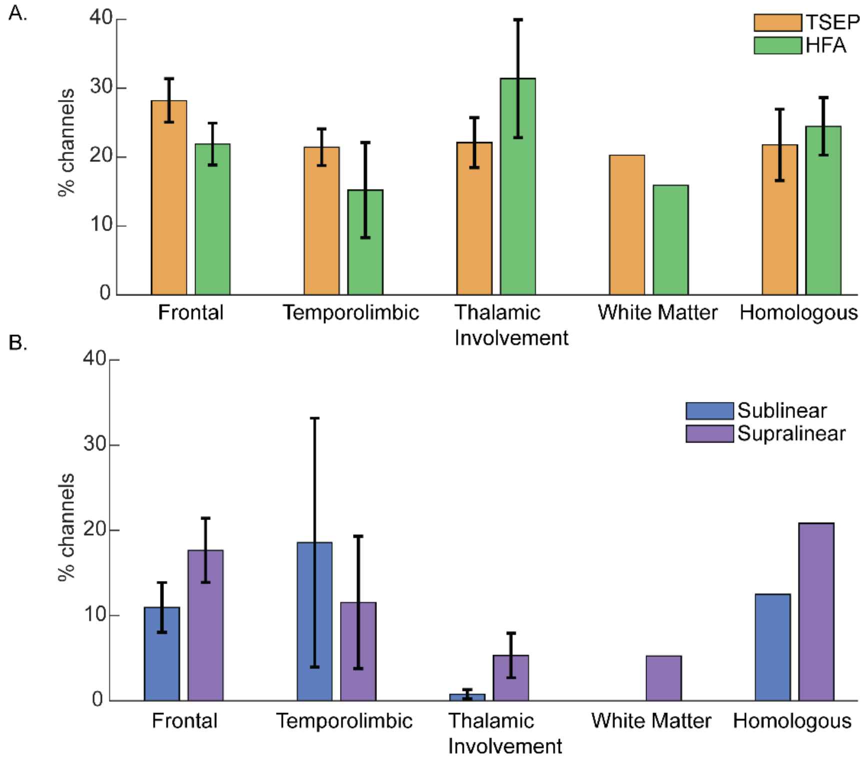


**Supplementary Figure 2:** Rates of additional responsiveness and nonlinearity as a function of stimulation location/cohort (7 mA). We observed a relatively even rate of TSEP additional responsiveness across the 5 stimulation cohorts, while nonlinearities appeared larger for frontal and temporolimbic cohorts compared to the thalamic involvement group. The low n (≤5) for cohorts other than frontal and thalamic involvement makes further statistical analysis infeasible.

**Table S2.1**

| TSEP detector | Detector thresholds | | | | TSEP rate (mean ± SEM %, n pairs) |
|---|---|---|---|---|---|
| | Minimum peak amplitude (z) | Minimum peak area (z·ms) | Minimum amplitude or area (× channel baseline) | First peak within (ms) | |
| Low threshold | 1.645 (p < 0.1) | 50 | 1.5 | 500 | 17.7 ± 1.2 (n = 87) |
| Moderate (used in the manuscript) | 1.96 (p < 0.05) | 100 | 2 | 500 | 20.4 ± 1.3 (n = 86) |
| High threshold | 2.576 (p < 0.01) | 200 | 3 | 300 | 25.7 ± 2.2 (n = 84) |
| Keller et al. (2011) | largest \|response\| at 50-500 ms > mean \|baseline\| + 6 SD (baseline -500 to -50 ms) | | | | 20.6 ± 1.5 (n = 84) |

**Table S2.2**

| HFA threshold | Thresholds | | HFA rate (mean ± SEM %, n pairs) |
|---|---|---|---|
| | Peak (× baseline power) | Area above 1 (s) | |
| Low | 2.0 | 0.10 | 22.4 ± 1.9 (n = 84) |
| Medium (used in the manuscript) | 2.5 | 0.15 | 25.1 ± 2.7 (n = 84) |
| High | 3.0 | 0.20 | 25.5 ± 2.8 (n = 81) |
| Very high | 4.0 | 0.25 | 25.8 ± 3.1 (n = 80) |
| Extra high | 5.0 | 0.30 | 24.7 ± 3.0 (n = 76) |

**Supplementary Table 2:** Sensitivity of additional responsiveness to classification threshold (7 mA).
The TSEP rate varies between 17.7%-25.7%, while the HFA rate is stable at 22.4-25.8% across the thresholds tested. The broader trend of 20-25% additional responsiveness to dual-site stimulation is robust to choice of threshold.

**Table S3**

| Minimum nonlinear period (ms) | Channels responsive to | Sublinearity rate (mean ± SEM %, n pairs) | Supralinearity rate (mean ± SEM %, n pairs) |
|---|---|---|---|
| 0 (any significant cluster) | both sites | 21.1 ± 2.6 (n = 63) | 24.0 ± 3.1 (n = 63) |
| | either site (A or B) | 10.3 ± 1.3 (n = 86) | 12.2 ± 1.6 (n = 86) |
| 50 | both sites | 10.0 ± 1.6 (n = 63) | 17.3 ± 2.4 (n = 63) |
| | either site (A or B) | 5.0 ± 0.7 (n = 86) | 8.4 ± 1.3 (n = 86) |
| 75 (used in the manuscript) | both sites | 7.5 ± 1.4 (n = 63) | 13.3 ± 2.1 (n = 63) |
| | either site (A or B) | 3.8 ± 0.6 (n = 86) | 5.9 ± 1.0 (n = 86) |
| 100 | both sites | 4.7 ± 1.1 (n = 63) | 8.7 ± 1.5 (n = 63) |
| | either site (A or B) | 2.7 ± 0.5 (n = 86) | 3.8 ± 0.7 (n = 86) |
| 150 | both sites | 2.8 ± 0.7 (n = 63) | 4.6 ± 1.1 (n = 63) |
| | either site (A or B) | 1.6 ± 0.3 (n = 86) | 2.0 ± 0.5 (n = 86) |

**Supplementary Table 3:**

Sensitivity of nonlinearity rate to classification threshold (0, 50, 75, 100, 150ms).

We observe that the absolute nonlinearity rate is sensitive to changes in the timing threshold for classification of a given channel as nonlinear. The use of a timing threshold acts to indirectly control the nonlinearity rate across channels. We chose the threshold of 75 ms since crossing that implies ≥10% of the response period was detected as nonlinear, but other choices could be argued to be similarly valid. However, the core trends we observe are applicable across thresholds - 1) channels responsive to both stimulation locations are twice as likely to be nonlinear than channels responsive to atleast one location. 2) supralinearities appear to be dominating sublinearities across thresholds at 7mA for channels responsive to stimulation at both locations. The timing histograms use timepoints across all channels, and are therefore invariant to choice of nonlinearity threshold.

**Table S4.1**

**Two-site stimulation: A&B on A+B (Figure 3c–e)**

| Amplitude: feature | Slope | Intercept | $R^2$ | p-value (slope) | n channels |
|---|---|---|---|---|---|
| 7 mA: Amplitude | 0.97 | 0.04 | 0.95 | $<10^{-300}$ | 4853 |
| 7 mA: AUC | 0.95 | 0.04 | 0.91 | $<10^{-300}$ | 4853 |
| 7 mA: Time to maximum peak | 0.46 | 89.11 | 0.25 | $<10^{-300}$ | 4848 |
| 4 mA: Amplitude | 1.01 | −0.18 | 0.95 | $<10^{-300}$ | 2543 |
| 4 mA: AUC | 1.02 | −0.02 | 0.89 | $<10^{-300}$ | 2543 |
| 4 mA: Time to maximum peak | 0.39 | 88.92 | 0.19 | $5.5\times10^{-118}$ | 2540 |
| 2 mA: Amplitude | 0.99 | −0.25 | 0.97 | $<10^{-300}$ | 574 |
| 2 mA: AUC | 0.90 | −0.01 | 0.88 | $3.7\times10^{-263}$ | 574 |
| 2 mA: Time to maximum peak | 0.54 | 72.37 | 0.29 | $3.0\times10^{-45}$ | 573 |

**Table S4.2**

**Three-site stimulation: A&B&C on A+B+C (Figure 5c)**

| Amplitude: feature | Slope | Intercept | $R^2$ | p-value (slope) | n channels |
|---|---|---|---|---|---|
| 7 mA: Amplitude | 0.93 | 0.03 | 0.94 | $<10^{-300}$ | 839 |
| 7 mA: AUC | 0.89 | 0.09 | 0.87 | $<10^{-300}$ | 839 |
| 4 mA: Amplitude | 0.96 | −0.16 | 0.95 | $<10^{-300}$ | 570 |
| 4 mA: AUC | 0.99 | −0.02 | 0.88 | $2.0\times10^{-265}$ | 570 |
| 2 mA: Amplitude | 0.96 | −0.30 | 0.96 | $8.5\times10^{-127}$ | 185 |
| 2 mA: AUC | 0.86 | 0.03 | 0.82 | $4.5\times10^{-69}$ | 185 |

**Supplementary Table 4:** Results of linear regression tests for plots of dual-site response vs sum of single site responses (2, 4, and 7 mA).

**Table S5.1. Linear mixed-effects model of Amp_AB (peak amplitude)**

| Term | β [95% CI] | Coef. p | Omnibus F ($df_1$) | Omnibus p |
|---|---|---|---|---|
| (Intercept) | 0.028 [−0.003, 0.059] | — | 3.21 (1) | 0.073 |
| **StimCohort** | | | **3.89 (5)** | **0.002** |
| Temporolimbic | 0.014 [−0.026, 0.054] | 0.498 | | |
| Homologous | −0.067 [−0.105, −0.030] | $4.1\times10^{-4}$ | | |
| White matter | −0.005 [−0.067, 0.057] | 0.873 | | |
| Thalamic involvement | −0.022 [−0.050, 0.005] | 0.116 | | |
| Other | −0.028 [−0.052, −0.003] | 0.027 | | |
| **anatomicRegion** | | | **1.64 (8)** | **0.109** |
| Temporal | 0.007 [−0.017, 0.031] | 0.585 | | |
| Central sulcus | −0.040 [−0.078, −0.003] | 0.037 | | |
| Parietal–occipital | −0.046 [−0.100, 0.009] | 0.101 | | |
| Limbic | −0.028 [−0.062, 0.007] | 0.119 | | |
| Insula | 0.019 [−0.021, 0.059] | 0.352 | | |
| Subcortical | −0.004 [−0.031, 0.023] | 0.788 | | |
| White matter | −0.003 [−0.019, 0.013] | 0.698 | | |
| Unknown | 0.015 [−0.014, 0.044] | 0.315 | | |
| isLocal | 0.008 [−0.010, 0.027] | — | 0.77 (1) | 0.381 |
| Amp_AplusB | 1.010 [0.995, 1.024] | — | $1.87\times10^{4}$ (1) | $<10^{-300}$ |
| AUC_AplusB | −0.045 [−0.059, −0.031] | — | 40.40 (1) | $2.3\times10^{-10}$ |
| AplusB_locMaxPk | −0.020 [−0.028, −0.012] | — | 22.46 (1) | $2.2\times10^{-6}$ |
| PeakOffset | −0.008 [−0.016, −0.000] | — | 4.03 (1) | 0.045 |

*Model: Amp_AB ~ Amp_AplusB + AUC_AplusB + AplusB_locMaxPk + PeakOffset + StimCohort + anatomicRegion + isLocal + (1|patientID)*

*Reference levels: StimCohort = Frontal, anatomicRegion = Frontal*

*Model fit: AIC = −1040.3, BIC = −904.1, log-likelihood = 541.2; n = 4851 observations from 20 participants (87 pairs)*

**Table S5.2. Linear mixed-effects model of AUC_AB (area under the curve)**

| Term | β [95% CI] | Coef. p | Omnibus F ($df_1$) | Omnibus p |
|---|---|---|---|---|
| (Intercept) | 0.007 [−0.042, 0.056] | — | 0.08 (1) | 0.772 |
| **StimCohort** | | | **6.05 (5)** | **$1.4\times10^{-5}$** |
| Temporolimbic | 0.065 [0.012, 0.119] | 0.016 | | |
| Homologous | −0.050 [−0.099, −0.000] | 0.048 | | |
| White matter | 0.063 [−0.019, 0.145] | 0.132 | | |
| Thalamic involvement | −0.045 [−0.082, −0.008] | 0.016 | | |
| Other | −0.012 [−0.045, 0.021] | 0.476 | | |
| **anatomicRegion** | | | **3.89 (8)** | **$1.4\times10^{-4}$** |
| Temporal | 0.029 [−0.002, 0.061] | 0.066 | | |
| Central sulcus | −0.087 [−0.137, −0.038] | $5.0\times10^{-4}$ | | |
| Parietal–occipital | −0.072 [−0.143, −0.001] | 0.046 | | |
| Limbic | −0.036 [−0.082, 0.009] | 0.115 | | |
| Insula | −0.011 [−0.062, 0.041] | 0.688 | | |
| Subcortical | 0.014 [−0.021, 0.049] | 0.444 | | |
| White matter | 0.017 [−0.004, 0.038] | 0.108 | | |
| Unknown | 0.007 [−0.030, 0.045] | 0.704 | | |
| isLocal | −0.018 [−0.042, 0.007] | — | 2.05 (1) | 0.152 |
| Amp_AplusB | 0.196 [0.177, 0.215] | — | 416.58 (1) | $6.9\times10^{-89}$ |
| AUC_AplusB | 0.775 [0.757, 0.793] | — | $7.10\times10^{3}$ (1) | $<10^{-300}$ |
| AplusB_locMaxPk | −0.023 [−0.034, −0.013] | — | 18.13 (1) | $2.1\times10^{-5}$ |
| PeakOffset | −0.004 [−0.014, 0.006] | — | 0.70 (1) | 0.404 |

*Model: AUC_AB ~ Amp_AplusB + AUC_AplusB + AplusB_locMaxPk + PeakOffset + StimCohort + anatomicRegion + isLocal + (1|patientID)*

*Reference levels: StimCohort = Frontal, anatomicRegion = Frontal*

*Model fit: AIC = 1516.4, BIC = 1652.7, log-likelihood = −737.2; n = 4851 observations from 20 participants (87 pairs)*

**Table S5.3. Linear mixed-effects model of AB_locMaxPk (time to maximum peak)**

| Term | β [95% CI] | Coef. p | Omnibus F ($df_1$) | Omnibus p |
|---|---|---|---|---|
| (Intercept) | −0.022 [−0.137, 0.093] | — | 0.14 (1) | 0.703 |
| **StimCohort** | | | **3.46 (5)** | **0.004** |
| Temporolimbic | 0.142 [−0.010, 0.293] | 0.066 | | |
| Homologous | 0.015 [−0.126, 0.156] | 0.833 | | |
| White matter | 0.040 [−0.195, 0.275] | 0.739 | | |
| Thalamic involvement | −0.110 [−0.215, −0.006] | 0.038 | | |
| Other | −0.053 [−0.146, 0.040] | 0.262 | | |
| **anatomicRegion** | | | **1.37 (8)** | **0.203** |
| Temporal | 0.055 [−0.036, 0.145] | 0.235 | | |
| Central sulcus | −0.018 [−0.160, 0.125] | 0.810 | | |
| Parietal–occipital | 0.209 [0.003, 0.414] | 0.047 | | |
| Limbic | 0.011 [−0.120, 0.143] | 0.865 | | |
| Insula | 0.023 [−0.127, 0.174] | 0.759 | | |
| Subcortical | 0.037 [−0.065, 0.139] | 0.473 | | |
| White matter | 0.046 [−0.014, 0.106] | 0.135 | | |
| Unknown | −0.087 [−0.196, 0.022] | 0.118 | | |
| isLocal | 0.083 [0.012, 0.154] | — | 5.21 (1) | 0.022 |
| Amp_AplusB | −0.078 [−0.133, −0.024] | — | 7.87 (1) | 0.005 |
| AUC_AplusB | 0.066 [0.013, 0.118] | — | 6.07 (1) | 0.014 |
| AplusB_locMaxPk | 0.616 [0.585, 0.648] | — | $1.51\times10^{3}$ (1) | $6.5\times10^{-287}$ |
| PeakOffset | 0.279 [0.249, 0.308] | — | 350.66 (1) | $1.4\times10^{-75}$ |

*Model: AB_locMaxPk ~ Amp_AplusB + AUC_AplusB + AplusB_locMaxPk + PeakOffset + StimCohort + anatomicRegion + isLocal + (1|patientID)*

*Reference levels: StimCohort = Frontal, anatomicRegion = Frontal*

*Model fit: AIC = 11844.4, BIC = 11980.6, log-likelihood = −5901.2; n = 4848 observations from 20 participants (87 pairs)*

**Supplementary Table 5:** Results of Linear Mixed Effect Models for two-site stimulation (7 mA).

**Table S6.1**

**Likelihood-ratio tests (all participants)**

| Response (Y) | n channels | logLik (1-feature) | logLik (complete) | $\chi^2$ (Δdf) | p | ΔAIC (complete − 1-feature) |
|---|---|---|---|---|---|---|
| Amp_AB | 4851 | 406.8 | 541.2 | 268.7 (18) | $1.2\times10^{-46}$ | −232.7 |
| AUC_AB | 4851 | −1106.8 | −737.2 | 739.2 (18) | $2.7\times10^{-145}$ | −703.2 |
| AB_locMaxPk | 4848 | −6183.9 | −5901.2 | 565.5 (18) | $1.7\times10^{-108}$ | −529.5 |

For every response the full model fits significantly better and has the lower AIC (negative ΔAIC favours the full model).

**Table S6.2**

**Leave-one-participant-out cross-validation ($R^2$ of the held-out predictions)**

| Response (Y) | n channels | 1-feature | Complete |
|---|---|---|---|
| Amp_AB | 4851 | 0.950 | 0.951 |
| AUC_AB | 4851 | 0.905 | 0.911 |
| AB_locMaxPk | 4848 | 0.242 | 0.292 |

*Each model was refitted with one participant left out and used to predict that participant's responses, once per participant; $R^2$ is computed over the held-out predictions of the n channels listed.*
*1-feature: Y ~ the same feature of the summed response (as the Figure 3c–e fits). Complete: the model of Supplementary Table 5.*

**Supplementary Table 6:** Likelihood-ratio tests comparing the full model against the single-feature model for each feature, and leave-one-participant-out cross-validation (7 mA)

### Table S7

| Term | β [95% CI] | Coef. p | Omnibus F ($df_1$) | Omnibus p |
|---|---|---|---|---|
| (Intercept) | −0.018 [−0.090, 0.054] | — | 0.24 (1) | 0.622 |
| **StimCohort** | | | **1.59 (2)** | **0.205** |
| Thalamic involvement | 0.049 [−0.021, 0.118] | 0.172 | | |
| Other | 0.011 [−0.046, 0.069] | 0.698 | | |
| **anatomicRegion** | | | **1.84 (8)** | **0.067** |
| Temporal | 0.057 [−0.002, 0.116] | 0.058 | | |
| Central sulcus | 0.043 [−0.066, 0.153] | 0.438 | | |
| Parietal–occipital | 0.046 [−0.081, 0.173] | 0.478 | | |
| Limbic | −0.026 [−0.103, 0.052] | 0.513 | | |
| Insula | 0.107 [0.010, 0.205] | 0.030 | | |
| Subcortical | 0.051 [−0.020, 0.122] | 0.159 | | |
| White matter | 0.010 [−0.033, 0.053] | 0.636 | | |
| Unknown | 0.101 [0.008, 0.193] | 0.034 | | |
| isLocal | −0.008 [−0.057, 0.041] | — | 0.10 (1) | 0.748 |
| Amp_AplusBplusC | 1.028 [0.990, 1.066] | — | $2.84\times10^{3}$ (1) | $4.4\times10^{-269}$ |
| AUC_AplusBplusC | −0.057 [−0.093, −0.021] | — | 9.76 (1) | 0.002 |
| AplusBplusC_locMaxPk | 0.001 [−0.024, 0.026] | — | 0.00 (1) | 0.951 |
| PeakOffset | 0.007 [−0.016, 0.031] | — | 0.37 (1) | 0.544 |

*Model: Amp_ABC ~ Amp_AplusBplusC + AUC_AplusBplusC + AplusBplusC_locMaxPk + PeakOffset + StimCohort + anatomicRegion + isLocal + (1|patientID)*

*Reference levels: StimCohort = Frontal, anatomicRegion = Frontal*

*Model fit: AIC = −45.1, BIC = 40.1, log-likelihood = 40.5; n = 839 observations from 7 participants (11 triplets)*

### Supplementary Table 7:

Results of Linear Mixed Effect Models for three-site stimulation (7 mA).

**Table S8.1**

| Cohort | TSEP additional responsiveness (mean ± SEM %, n pairs) | HFA additional responsiveness (mean ± SEM %, n pairs) |
|---|---|---|
| Frontal | 28.2 ± 3.1 (n = 14) | 21.9 ± 3.0 (n = 14) |
| Temporolimbic | 21.4 ± 2.6 (n = 5) | 15.2 ± 6.9 (n = 5) |
| Thalamic involvement | 22.1 ± 3.6 (n = 14) | 31.4 ± 8.6 (n = 14) |
| White matter | 20.3 (n = 1) | 15.9 (n = 1) |
| Homologous | 21.8 ± 5.2 (n = 5) | 24.5 ± 4.2 (n = 5) |
| Other | 17.3 ± 1.6 (n = 47) | 25.4 ± 4.2 (n = 45) |

**Table S8.2**

| Cohort (channels responsive to both sites) | Sublinearity rate (mean ± SEM %, n pairs) | Supralinearity rate (mean ± SEM %, n pairs) |
|---|---|---|
| Frontal | 11.0 ± 2.9 (n = 13) | 17.7 ± 3.8 (n = 13) |
| Temporolimbic | 18.6 ± 14.6 (n = 3) | 11.5 ± 7.8 (n = 3) |
| Thalamic involvement | 0.8 ± 0.5 (n = 11) | 5.3 ± 2.6 (n = 11) |
| White matter | 0.0 (n = 1) | 5.3 (n = 1) |
| Homologous | 12.5 (n = 1) | 20.8 (n = 1) |
| Other | 7.4 ± 1.8 (n = 34) | 14.4 ± 3.3 (n = 34) |

**Table S8.3**

| Cohort (channels responsive to either site (A or B)) | Sublinearity rate (mean ± SEM %, n pairs) | Supralinearity rate (mean ± SEM %, n pairs) |
|---|---|---|
| Frontal | 7.3 ± 1.8 (n = 14) | 11.1 ± 3.0 (n = 14) |
| Temporolimbic | 6.7 ± 5.7 (n = 5) | 5.0 ± 4.3 (n = 5) |
| Thalamic involvement | 0.9 ± 0.4 (n = 14) | 1.9 ± 1.1 (n = 14) |
| White matter | 0.0 (n = 1) | 2.9 (n = 1) |
| Homologous | 3.1 ± 1.5 (n = 5) | 4.6 ± 2.2 (n = 5) |
| Other | 3.5 ± 0.7 (n = 47) | 5.8 ± 1.5 (n = 47) |

"Other" consists of channels which have the stimulation sites in different lobes, such as frontal and temporal lobe, or frontal lobe in different hemispheres (which are not homologous areas), or one is in white matter while the other is in cortex, if they are in other subcortical structures, or marked unknown by parcellation.

**Supplementary Table 8:** Additional responsiveness and nonlinearity by stimulation cohort (7 mA).

**Table S9**

| Amplitude | Measure | Three-site (mean ± SEM %, n triplets) | Two-site, constituent pairs (mean ± SEM %, n pairs) | Unpaired test (three-site − two-site) | | |
|---|---|---|---|---|---|---|
| | | | | Difference (%) | p | Cliff's δ |
| 2 mA | TSEP added | 40.0 ± 14.7 (n = 6) | 31.0 ± 6.5 (n = 17) | 9.1 | 0.517 | 0.12 |
| | HFA added | insufficient (n = 3) | 28.4 ± 13.1 (n = 10) | not tested (fewer than 5 rates in a group) | | |
| | Sublinear | insufficient (n = 0) | insufficient (n = 2) | not tested (fewer than 5 rates in a group) | | |
| | Supralinear | insufficient (n = 0) | insufficient (n = 2) | not tested (fewer than 5 rates in a group) | | |
| 4 mA | TSEP added | 30.9 ± 6.2 (n = 11) | 29.2 ± 4.5 (n = 32) | 1.6 | 0.864 | 0.09 |
| | HFA added | 31.1 ± 6.3 (n = 11) | 31.8 ± 6.5 (n = 31) | −0.7 | 0.958 | 0.17 |
| | Sublinear | 7.0 ± 4.3 (n = 5) | 5.8 ± 2.8 (n = 17) | 1.2 | 0.887 | 0.09 |
| | Supralinear | 17.9 ± 8.5 (n = 5) | 5.5 ± 2.0 (n = 17) | 12.5 | 0.034 | 0.40 |
| 7 mA | TSEP added | 23.7 ± 5.0 (n = 11) | 20.8 ± 2.3 (n = 33) | 2.9 | 0.572 | 0.05 |
| | HFA added | 34.4 ± 6.3 (n = 11) | 31.1 ± 4.6 (n = 33) | 3.3 | 0.716 | 0.17 |
| | Sublinear | 14.6 ± 4.2 (n = 9) | 7.5 ± 1.7 (n = 28) | 7.0 | 0.070 | 0.33 |
| | Supralinear | 19.7 ± 8.2 (n = 9) | 10.5 ± 2.3 (n = 28) | 9.2 | 0.139 | 0.13 |

**Supplementary Table 9:** Additional responsiveness and nonlinearity with three-site and two-site stimulation (Figure 5b; 2, 4 and 7 mA).
